# The open to closed D-loop conformational switch determines length in filopodia-like actin bundles

**DOI:** 10.1101/2024.08.09.607305

**Authors:** Jonathan R Gadsby, Pantelis Savvas Ioannou, Richard Butler, Julia Mason, Alison Smith, Ulrich Dobramysl, Stacey Chin, Claire Dobson, Jennifer L Gallop

**Affiliations:** Gurdon Institute, University of Cambridge, Tennis Court Road, Cambridge, UK, CB2 1QN; Department of Biochemistry, University of Cambridge, Tennis Court Road, Cambridge, UK, CB2 1GA; Biologics Engineering, Oncology R&D, AstraZeneca, Cambridge, UK; Discovery Sciences, BioPharmaceuticals R&D, AstraZeneca, Cambridge, UK; Peter Medawar Building for Pathogen Research, Nuffield Department of Medicine, University of Oxford, Oxford, UK

## Abstract

Filopodia, microspikes and cytonemes are implicated in sensing the environment and in dissemination of morphogens, organelles and pathogens across tissues. Their major structural component is parallel bundles of actin filaments that assemble from the cell membrane. Whilst the length of filopodia is central to their function, it is not known how their lengths are determined by actin bundle dynamics. Here, we identified a set of monoclonal antibodies that lengthen filopodia-like structures formed in a cell-free reconstitution system, and used them to uncover a key molecular switch governing length regulation. Using immunolabelling, enzyme-linked immunosorbent assays, immunoprecipitation and immunoblock experiments, we identified four antibodies that lengthen actin bundles by selectively binding the open DNase 1-binding loop (D-loop) of actin filaments. The antibodies inhibit actin disassembly and their effects can be alleviated by providing additional actin or cofilin. This work indicates that maintaining an open state of the actin filament D-loop is a mechanism of generating the long projections of filopodia-like actin bundles.

## Introduction

Filopodia are composed of tightly bundled parallel F-actin filaments that form the core of thin finger-like protrusions from the plasma membrane. They have multiple roles in cell differentiation, migration, pathogen infection and environment sensing [1,2]. As exploratory organelles, the stochastic protrusion and retraction of filopodia results in structures of varying lengths that can reach from the cell body to the extracellular matrix, to other cells and across tissues in the case of cytonemes [3,4]. Filopodia lengths are typically up to 10 microns, while cytonemes can reach 10 times further. Growth or shrinkage rates are typically 1-2 µm/min and reaching ∼25 µm/min [5]. The underlying mechanisms determining filopodial length will involve balancing the rates of actin incorporation, bundling, severing and disassembly with lifetime of each phase [6]. What makes any given filopodium long or short remains an open question, yet is crucial for understanding the molecular basis of filopodial formation and how filopodia respond to intracellular and extracellular signals that guide cell sensing and movement.

Candidate determinants of filopodia length include include F-actin conformational changes, as well as the influence of actin binding proteins formins [7–9], myosin-X [10,11], fascin [12,13], capping protein [14,15], and cofilin [16,17]. Reconstitutions of fascin-mediated bundles emerging from Arp2/3 complex-nucleated actin with either VASP or mDia2 showed that mDia2 gives longer filopodia than VASP [8]. While myosin-X overexpression results in elongated filopodia, this depends on the tail domain rather than the motor domain [11]. Phosphorylation site mutants of fascin affect filopodial lengths [12]. In native neuronal filopodia cofilin-bound actin filaments are present in longer filopodia and are more common towards the cell body, consistent with increased affinity of F-actin for cofilin that occurs as the actin filament ages [16–18]. When actin polymerizes, G-actin monomers are initially incorporated into filaments with ATP occupying the actin nucleotide binding pocket. Filament formation causes rapid ATP hydrolysis. In the ADP-Pi state, hydrolysis has occurred but inorganic phosphate remains within the nucleotide binding pocket before it is slowly released, eventually leaving older filaments in the less stable ADP-bound state [18–20]. Along a filament there is a nucleotide gradient which locally marks its age, and this shifts the conformation of actin monomers and mechanical properties of the filament [21–23], including at a flexible site on the filament surface, the DNase 1-binding loop (D-loop). Cofilin binds to the closed D-loop conformation of ADP-actin filaments causing a twist in the filament that increases the depolymerization rate from the pointed end and diassembles actin filaments [24–27]. Cofilin localises as cofilactin filaments at the base of long filopodia in neurons where it may or may not be the trigger for actin depolymerization.

To explore the molecular determinants of filopodial length regulation in an unbiased way we conducted a phenotypic antibody screen on a cell-free system of filopodia-like structures (FLS). FLS are nucleated on addition of *Xenopus* egg high-speed supernatant extracts (HSS) to PI(4,5)P_2_-containing supported lipid bilayers [28]. FLS are comprised of fascin-bundled parallel actin filaments and recapitulate the growth and shrinkage dynamics seen in cellular filopodia [5]. We previously used FLS in a phenotypic screen as antigen for phage display selections with a library of human single chain variable fragments (scFvs) to identify antibodies that specifically bound FLS. After incubation of the *Xenopus* egg extracts with the identified scFvs, FLS were grown and their morphological properties analysed, leading to the isolation of antibodies that cause longer and shorter FLS [29]. Nanobody reagents have successfully isolated against fascin previously [30,31] and the difference with our FLS approach is the ability to identify novel antigens.

Here we have identified the antigen and epitope of several antibodies that make FLS longer and determined their mechanism of action. Using immunostaining, enzyme-linked immunosorbent assays and displacement of actin-binding proteins, we show that four independent antibody clones recognise the open DNase1-binding loop conformation of actin filaments. By measuring FLS length and actin assembly/disassembly rates using photobleaching, we show that the disassembly rate is diminished by antibody binding while the assembly rate is unchanged. The antibodies’ effect on lengthening FLS can be overcome by increasing cofilin and actin concentrations in the extracts. We propose that a switch in actin filament D-loop conformation from the open to closed state within the actin bundle is an underlying molecular feature of length control in filopodia and related actin networks.

## Results

### A group of mAbs identified by phage display phenotypic screening lengthens FLS actin bundles

Incubation of HSS with a supported lipid bilayer enriched in PI(4,5)P_2_ yields the assembly of FLS that are readily followed by spinning disk confocal microscopy z-stacks followed by 3D reconstruction and image-based quantification using our pipeline FLSAce [5,28]. We converted the scFvs selected in our previous phage display phenotypic screen on FLS [29] into bivalent full antibodies expressing them as HuIgG1s and preincubated them with HSS before addition to the supported lipid bilayer. The FLS were allowed to grow and imaged after 25-35 minutes when the balance of growth and shrinkage results in a steady-state. Incubation with four antibodies: mAb2, mAb6, mAb14 and mAb17 resulted in FLS of significantly longer lengths, similar to their scFv counterparts, whereas the isotype control antibody R347 was similar to the untreated control and had no effect on FLS (Fig 1A-B).

**Fig 1.**
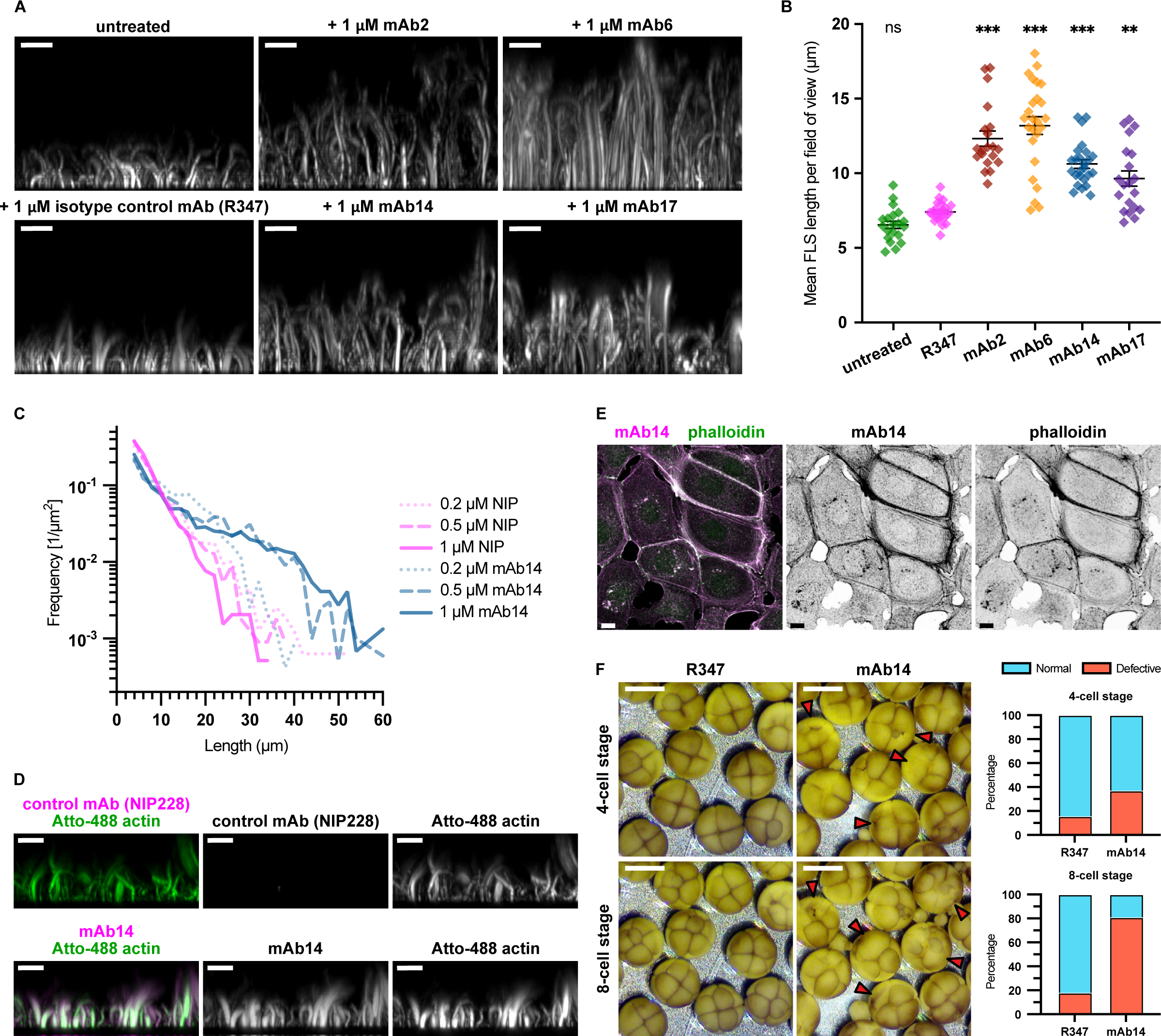
A group of antibodies lengthen FLS and immunolabel along F-actin structures. **(A)** Each of mAb2, 6, 14 and 17 produces FLS of significantly longer length compared to the control R347 antibody, which is comparable to untreated FLS. Images show maximum intensity projections of x-z planes from a spinning disc confocal z-stack of Atto-488-actin in representative FLS regions. Scale bars = 10 μm. **(B)** Quantification of mean FLS length per field of view in each condition. N=25 (R347, mAb6, mAb14) or 20 (mAb2, mAb17) fields of view. Lines indicate mean ± SEM. Significance was assessed by ordinary one-way ANOVA with Sidak’s multiple comparison test (R347 vs UNT: p=0.539 [ns], R347 vs mAb2, mAb6 or mAb14: all p<0.001 [***], R347 vs mAb17: p=0.001 [**]). **(C)** Log frequency against FLS length histogram showing the dose responsive shift in FLS length distribution on mAb14 treatment. n(FLS) = 1579 (0.2 µM NIP), 1105 (0.5 µM NIP), 1944 (1 µM NIP), 2394 (0.2 µM mAb14), 1997 (0.5 µM mAb14), & 1464 (1 µM mAb14). **(D)** mAb14 labels along the length of the FLS, while the NIP228 control does not. Maximum intensity projections of x-z planes of Atto-488-actin (green) in representative untreated FLS regions immunostained for either the isotype control mAb NIP228 or mAb14 (magenta). Scale bars = 10 μm. **(E)** mAb14 (magenta) labels F-actin in HEK293T cells similarly to Alexa-488-phalloidin marked by the phalloidin stain (green). Maximum intensity projections of x-y plane of a laser scanning confocal z-stack. Scale bars = 10 μm. **(F)** Images of cleaving *Xenopus* embryos at the 4 and 8 cell stage showing that mAb14 injected cells have delayed cytokinesis compared to the uninjected half or R347 control injected cells. Graphs illustrate scoring of each embryo as to whether cytokinesis appears normal or defective at each stage. n(embryos): 61 (R347) and 58 (mAb14). Scale bars = 1 mm.

As we previously observed, the lengths of unperturbed FLS in a stationary state of growth can be well-described by an exponential distribution, which is intrinsically linked to their growth dynamics [5]. Plotting these distributions on a log-scaled y-axis allowed the nature of the length change driven by the antibodies to be discerned (Fig 1C): in the unperturbed control condition, the number of FLS decreases sharply with increasing length, with no FLS longer than roughly 32 µm detectable; in contrast, the lengthening antibodies (mAb2, 6, 14, 17) instead cause a considerable increase of the number of FLS at longer lengths, altering the shape of the distribution (Fig 1C). In reconstitutions of filopodia-like networks from purified proteins, high concentrations of formins also lengthened the resulting actin bundles changing the length distribution in a similar way [8].

The change in FLS length arose in a dose-responsive manner for each antibody, while there was no change with either isotype control antibody R347 and NIP228 (Fig 1C, S1). To begin to identify the antigen of the antibodies, we used immunostaining and Western blotting. While by immunostaining the four lengthening mAbs were capable of staining the full length of the actin shaft of untreated FLS (Fig 1D), no band was obtained using the antibodies for Western blotting of HSS (unpublished observations). No staining of the FLS shaft was observed with either the isotype control antibody NIP228 or mAb4 from the same phenotypic screen (Fig1D, S2A), which we had previously identified as an anti-SNX9 antibody [29]. To test whether a similar staining pattern was observed in cells, we immunostained HEK293 cells together with Alexa-488 conjugated phalloidin for filamentous actin, seeing close correspondence, which was confirmed by controls (Fig 1E, S2B). Microinjection of mAb14 into one cell of *Xenopus* embryos at the 2 cell stage led to defective formation of the cytokinetic furrow in the injected cell compared with R347 control antibody, consistent with a disruption in F-actin (Fig 1F).

### The group of FLS lengthening antibodies specifically recognise F-actin

Next, we performed immunoprecipitation experiments with mAbs 2, 6, 14 and 17 in the presence and absence of actin monomer sequestering drug latrunculin B [32]. A distinct band of approximately 45 kDa was isolated from HSS by each of mAb 2, 6, 14 and 17 and not the R347 isotype control, which was prevented by the latrunculin B treatment (Fig 2Ai, full gels are shown in Fig S3). A subsequent Western blot using a commercial pan-actin antibody identified the immunoprecipitated protein as actin (Fig 2Aii). To determine whether purified actin was sufficient to bind the antibodies we performed sedimentation experiments with purified rabbit muscle actin under conditions of ∼50% G/F-actin, with a high speed sedimentation to separate F-actin filaments and low speed sedimentation to separate actin bundles (Fig 2B-C). The R347 control antibody remains predominantly in the supernatant with G-actin. In contrast, each of mAb 2, 6, 14 and 17 sedimented in the pellet with F-actin filaments and F-actin bundles. There was some increase in actin sedimenting in the pellet (e.g. mAbs 6 and 17), our conclusion from trial experiments is that it originates from variable disruption of fragile pellets (Fig 2B-C).

**Fig 2.**
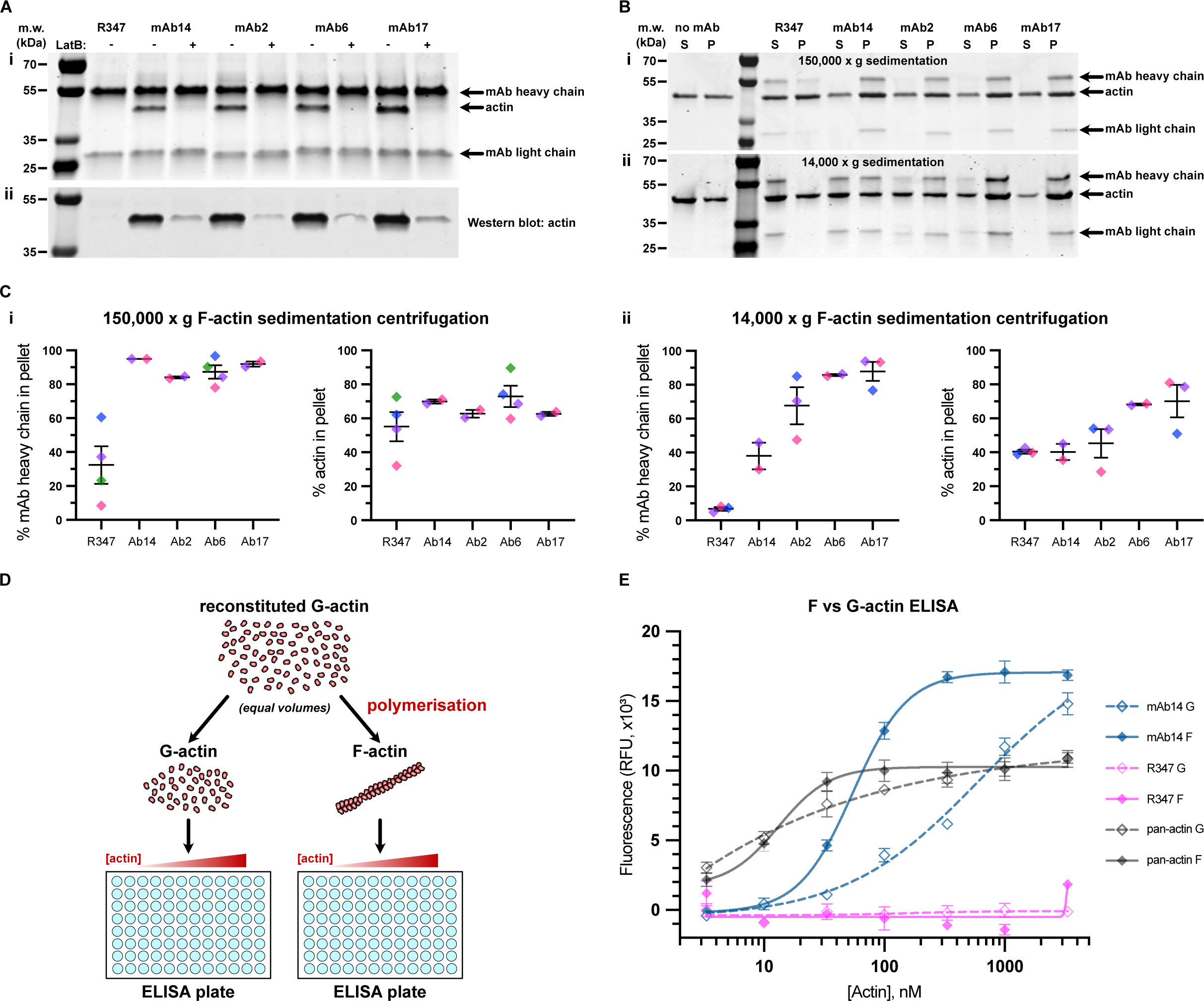
FLS lengthening mAbs specifically recognise, and shift the equilibrium towards bundled F-actin. **(A)** mAbs 14, 2, 6, and 17 co-immunoprecipitate a 45 kDa protein, visible by **(i)** Coomassie staining, that is prevented in the presence of actin monomer sequestering drug latruculin B (LatB). **(ii)** Anti-pan-actin Western blot of the Co-IP shown in **(i)**, indicating that the 45 kDa band observed is actin. **(B-C) (i)** mAbs 2, 6, 14 and 17 sediment with purified F-actin at 150,000*g*. Coomassie gel **(B)** and densitometry analysis **(C)** showing G-actin in the supernatant (S) and F-actin in the pellet (P). with a slight increase in pelleted actin with the antibodies. **(ii)** mAbs 2, 6, 14 and 17 sediment with purified F-actin at 14,000*g*. Coomassie gel **(B)** and densitometry analysis **(C)** showing G-actin in the supernatant (S) and bundled F-actin in the pellet (P) with a slight increase in pelleted actin with the antibodies. The R347 control mAb remains in supernatant. N-2-4 assays per condition. Lines indicate mean ± SEM, colours indicate datapoints from the same assay. **(D)** Schematic illustrating the experimental set-up of an ELISA designed to test the specificity of mAb to G or F-actin. An equal dilution series of G or F-actin is bound to an ELISA plate and probed with each mAb of interest. **(E)** Response fitted binding curves for mAb14, R347 and a pan-actin antibody binding to F or G-actin. Symbols and error bars indicate the mean and SEM of each measured concentration (each data point was taken in triplicate). mAb14 binds to F-actin with greater affinity than G-actin, in contrast to the pan-actin antibody and the R347 isotype control (which does not bind).

To confirm the antibody specificity for F-actin we designed an actin-down ELISA. Reconstituted rabbit skeletal muscle actin was split into two equal volumes: one half of which was prepared as monomeric G-actin, with the other half polymerized into F-actin filaments (Fig 2D). The two samples were prepared into a dilution series on an ELISA plate, and probed with mAbs 2, 6, 14 and 17, the R347 isotype control, and a pan-actin primary antibody that is able to detect both G and F-actin. As expected, R347 failed to bind, and we verified that the pan-actin control bound to G and F-actin similarly. This shows that the locally increased concentration achieved by clustering of actin monomers into filaments does not necessarily increase antibody binding by avidity effects. Each of the lengthening antibodies bound F-actin in preference to G-actin (Fig 2E, S3E). Quantification of the EC50 values from the binding curves suggested an approximately 10-fold increase in affinity of mAb 2, 6, 14 and 17 towards F-actin relative to G-actin (Fig S3F).

### Epitope mapping using the displacement of the actin-binding probes

To test whether the antibodies were capable of displacing known actin binding proteins we used GFP-utrophin tandem CH1/CH2 domains (UtrCH) [33], which normally labels the full length of FLS, and observed that the probe was displaced from FLS in the presence of each of mAb 2, 6, 14 and 17 (Fig 3A, S4A). At the 1 µM antibody concentration, the GFP-UtrCH was restricted to a thin strip nearest the supported lipid bilayer (Fig 3A). The displacement of GFP-UtrCH and the increase in length of FLS, were similarly dose-responsive for each mAb (Fig 3A-B, S4B-C). To quantify the displacement, we measured the mean fluorescence contributed by all FLS over the field of view at each z-slice for both GFP-UtrCH and Alexa-568 actin channels, showing that the extent of GFP-UtrCH is reduced by higher concentrations of antibody and that only fluorescent actin allows the visualization of the longer FLS length in the presence of mAb14 (Fig 3C, S4A-D).

**Fig 3.**
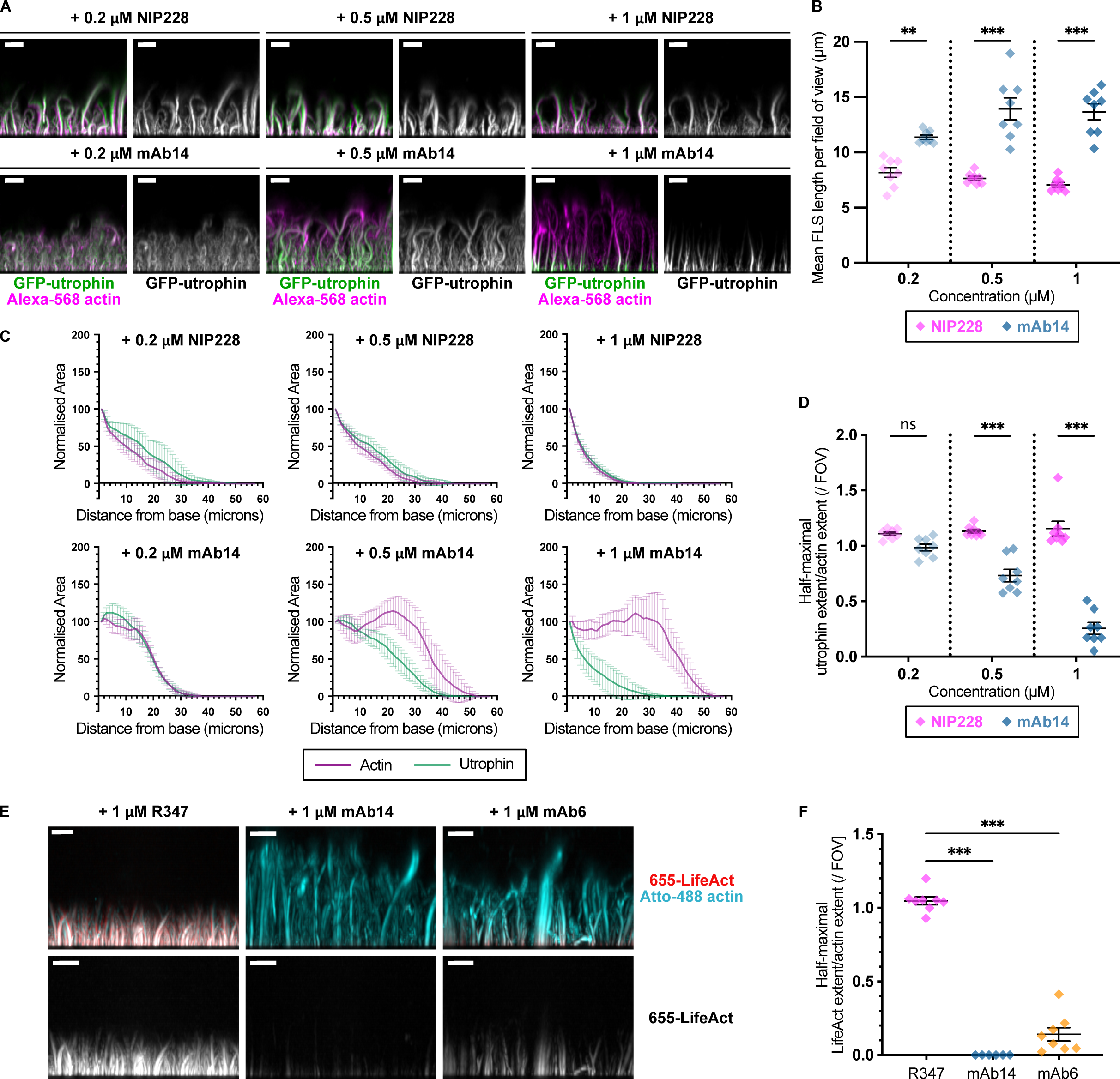
FLS lengthening mAbs displace the actin binding probes GFP-UtrCH and LifeAct from FLS actin bundles. **(A)** mAb14 prevents the binding of GFP-UtrCH to FLS in a dose-responsive manner, while the GFP-UtrCH stains similarly to Alexa568-actin on treatment with the NIP228 antibody control. Maximum intensity projections of x-z plane of a spinning disc confocal z-stack. All scale bars = 10 μm. **(B)** mAb14 increases path length and NIP228 does not. Mean FLS length per field of view from N=8 fields of view. Lines indicate mean ± SEM. Significance was assessed by ordinary one-way ANOVA with Sidak’s multiple comparison test: 0.2 µM NIP228 vs 0.2 µM mAb14, p = 0.002 [**], 0.5 µM NIP228 vs 0.5 µM mAb14 & 1 µM NIP228 vs 1 µM mAb14, p < 0.001 [***]. Further comparisons can be found in Table S1. **(C)** Quantification of GFP-UtrCH labelling compared with Alexa-568 actin for each condition. There is no difference with different concentrations of NIP228, whilst UtrCH is displaced on the addition of mAb14 in a dose responsive manner. Normalised mean actin and GFP-Utr fluorescent areas at each z plane for N=8 imaging regions per condition. Error bars indicate 95% confidence intervals. **(D)** Summary data from quantifications in C. Ratios of half maximal UtrCH fluorescence compared to the actin signal. Lines indicate mean ± SEM. Significance was assessed by ordinary one-way ANOVA with Sidak’s multiple comparison test: 0.2 µM NIP228 vs 0.2 µM mAb14, p = 0.456 [ns], 0.5 µM NIP228 vs 0.5 µM mAb14 & 1 µM NIP228 vs 1 µM mAb14, p < 0.001 [***]. Further comparisons can be found in Table S1. **(E)** LifeAct is also displaced by the lengthening antibodies. Maximum intensity projections of x-z plane of a spinning disk confocal z-stack showing displacement of LifeAct (red) compared with actin (cyan) by mAb14 and mAb6 in representative FLS regions preincubated with 1 μM R347, mAb6 or mAb14. All scale bars = 10 μm. **(F)** Quantification of LifeAct extent in each condition. There is no difference with different concentrations of NIP228, whilst LifeAct is strongly displaced by mAb6 and almost abolished by mAb14. Each datapoint represents an individual imaging region. N=6-8 fields of view per condition. Lines indicate mean ± SEM. Significance was assessed by ordinary one-way ANOVA with Sidak’s multiple comparison test: for both R347 vs mAb14 & R347 vs mAb6, p < 0.001 [***].

We noted that the GFP-UtrCH binds to an area of F-actin including the D-loop [22,34,35]. To verify that the lengthening antibodies interact with the D-loop we tested a second probe, LifeAct, a 17 residue peptide from the yeast protein Abp140, that has a smaller footprint on the F-actin filament more closely localized to the D-loop region (Fig 3E). Atto-655-LifeAct is displaced in the presence of either 1 µM mAb14, or mAb6 while the Atto-488-actin was similar to FLS treated with the control mAb (Fig 3E-F, S4F). The degree of LifeAct displacement was exacerbated compared to GFP-UtrCH, in particular for mAb14, suggesting that the D-loop is the predominant site of antibody binding (Fig 3E-F).

We wondered how common it is for anti-actin antibodies to cause a lengthening of actin bundles and displacement of actin-binding probes. We tested a panel of monoclonal anti-actin antibodies for these properties finding that most do not lengthen FLS or displace GFP-UtrCH (Fig S5). An antibody raised against residues 29-39 of human actin, which is adjacent to the D-loop, caused a slight lengthening effect at the low concentration attainable (Fig S5). This suggests that the D-loop is an epitope specific to the FLS lengthening antibodies, and that D-loop binding causes increased FLS length.

### The group of FLS lengthening antibodies specifically recognise F-actin in the open D-loop conformation

Recent studies have shown that the D-loop at the surface of filaments undergoes conformational changes linked to these different nucleotide states [34,36]. The D-loop is the most flexible region at the filament surface, and inserts into a hydrophobic cavity between the SD1 and SD3 domains of the neighbouring N^+2^ monomer. The D-loop switches between an “open” and “closed” conformation, with ATP and ADP-Pi nucleotide states containing a mix of the two conformations whilst ADP bound filaments are only found in the closed state [21]. The D-loop is a key site for actin regulatory protein binding, with different effectors recognising different nucleotide and conformational states; for example, the depolymerization factor cofilin recognises ADP-actin in the closed conformation, whereas coronin strongly prefers the ADP-Pi state [21,22,27,37].

Phalloidin and jasplakinolide are two natural actin stabilizing compounds that compete for binding at the same site (away from the D-loop) within the actin filament [38,39]. Importantly, they can be used to stabilize polymerized actin in contrasting D-loop conformations allosterically. Aged ADP F-actin is stabilized in the closed conformation by phalloidin, whereas jasplakinolide is able to switch the D-loop back to an open conformation (despite the absence of the released inorganic phosphate) [21,39–41]. We therefore modified our F-actin binding ELISA, using polymerized filaments stabilized with either jasplakinolide or phalloidin to determine the binding affinity of the antibodies to specifically open vs closed actin D-loop conformations. mAb14 bound to both stabilized groups with greater affinity than untreated F-actin with a strong preference for jasplakinolide-stablized “open” conformation filaments over phalloidin-treated “closed” conformation F-actin (Fig 4A). In contrast, the pan-actin control antibody which recognises F- and G-actin with the same specificity showed no such preference, and in fact treatment with either compound compromised binding (Fig 4B). mAbs 2, 6 and 17 all showed a similar result to mAb14, with in each case an approximate 10-fold greater EC50 affinity towards jasplakinolide over phalloidin-treated F-actin (Fig S6). We wondered whether the jasplakinolide or phalloidin stabilization of filaments would affect FLS length similar to the the antibodies. Treatment with either compound resulted in a modest decrease in FLS length and fuzzy morphology consistent with native actin filament conformations being an important aspect of correct FLS morphology (Fig S7).

**Fig 4.**
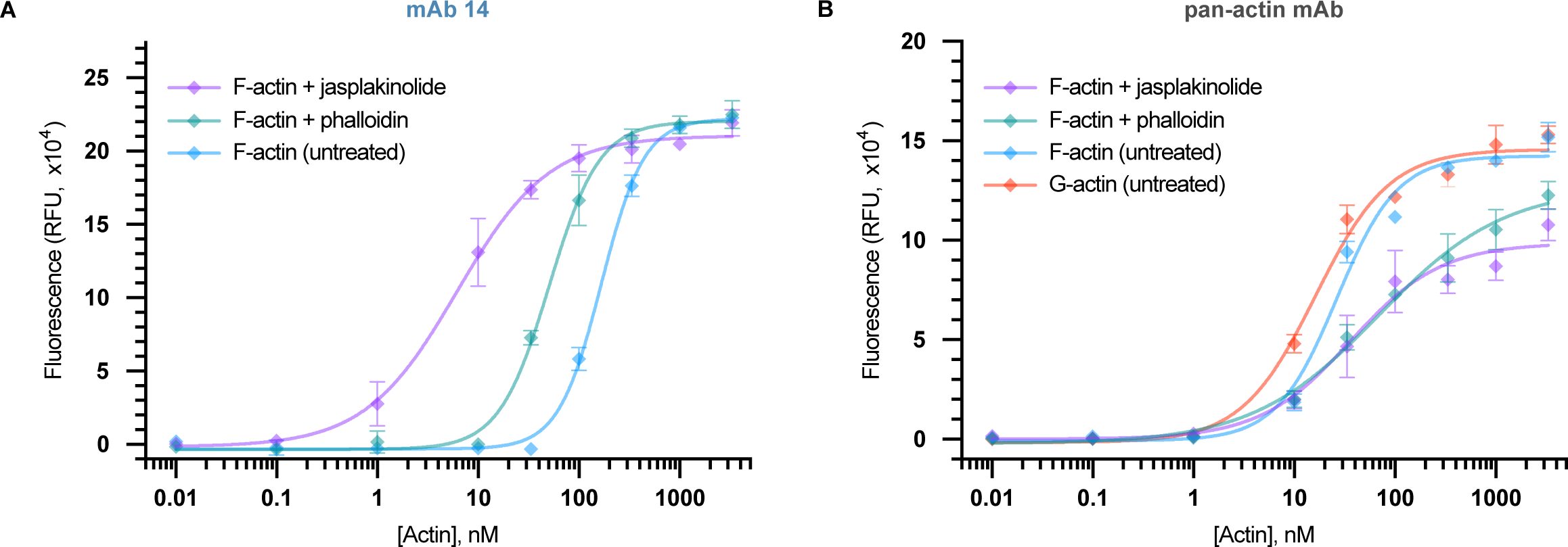
FLS lengthening mAbs specifically recognise the open conformation of the actin D-Loop. **(A)** mAb14 binds the open conformation stabilised by jasplakinolide and or closed phalloidin-treated F-actin conformation better than aged ADP:actin. Symbols and error bars indicate the mean and SEM of each measured concentration (each data point was taken in triplicate). **(B)** Pan-actin antibody recognises F and G actin similarly with a slightly higher affinity than either phalloidin or jasplakinolide-treated actin. Symbols and error bars indicate the mean and SEM of each measured concentration (each data point was taken in triplicate).

### FLS lengthening antibodies protect actin filaments from disassembly

The lengthening effect of the antibodies on FLS implies that their interaction with the open D-loop conformation will act to either increase actin assembly or decrease disassembly. To test this, FLS were initiated and allowed to grow for either 5 minutes (to capture structures in the initial growing phase), or 25 minutes (when FLS have typically reached steady-state with growth balanced by shrinkage). The fluorescently-labelled actin in the entire field of view was then bleached and imaged every minute for the next 10 minutes to observe the reincorporation of bright freshly-added actin assembling at FLS tips and disassembly of faintly-fluorescent bleached FLS (Fig 5A).

**Fig 5.**
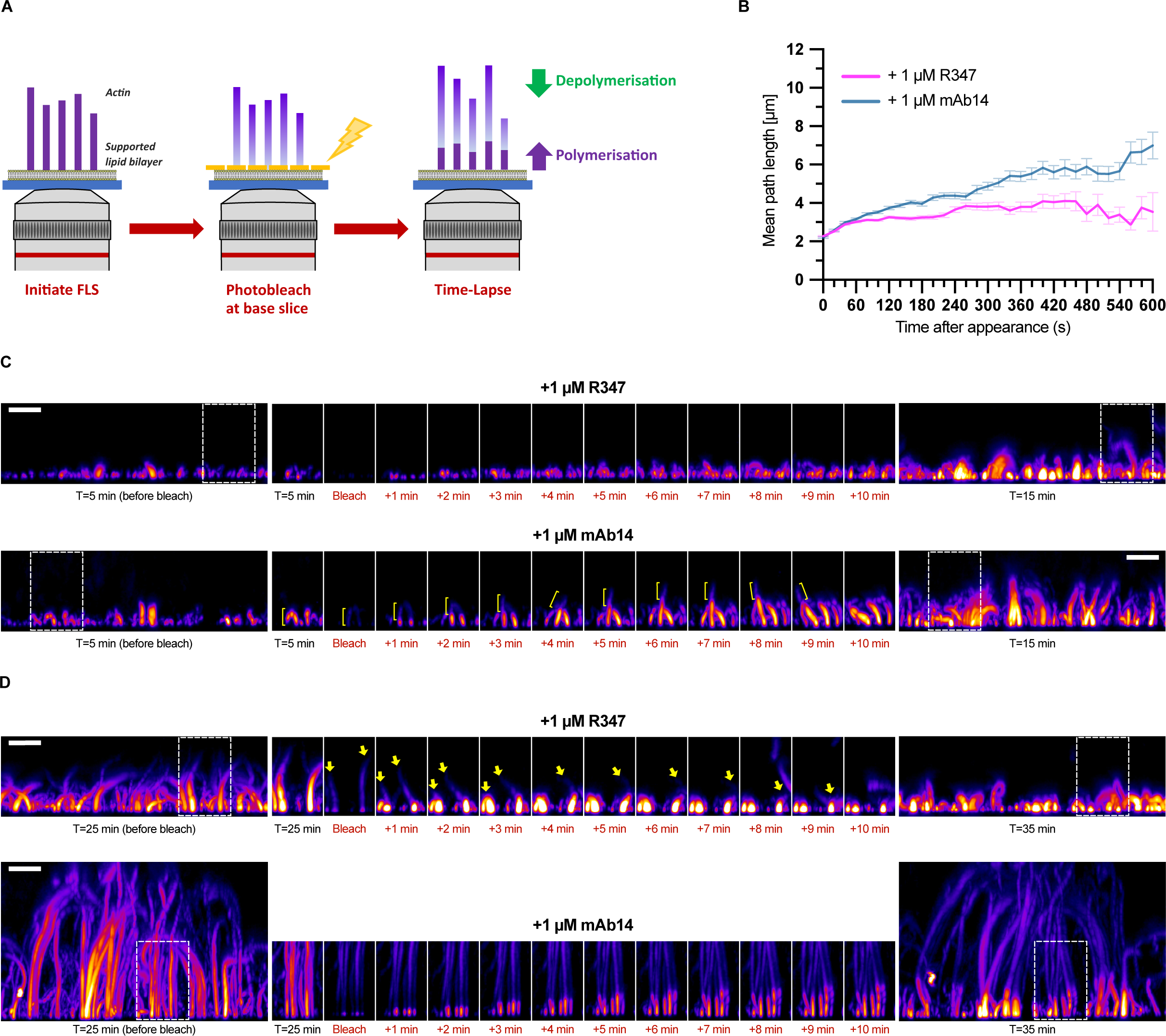
Lengthening mAbs protect FLS from depolymerization. **(A)** Schematic outlining the photobleaching experimental set-up used to examine FLS dynamics. FLS are allowed to grow for 5 minutes (to capture the initial growth phase) or 25 minutes (to reach steady state). The fluorescently tagged actin incorporated into FLS was then photobleached, and a time course taken every 20s for a further 10 minute period. The bleached actin present at the start of the time-lapse is readily distinguishable from the much brighter freshly incorporated actin, allowing both the incorporation rates of fresh actin, and the behaviour of the older bleached actin to be followed. **(B-C)** FLS treated with either R347 or mAb14 show similar initial rates of fresh actin incorporation at 5-15 min timepoints, however older bleached actin is resistant to disassembly with mAb14 treatment leading to growing disparity in length over time compared to R347. **(B)** Profile of mean FLS lengths at each time point relative to first detection. n(FLS) = 1417 (R347) and 981 (mAb14) combined from five imaging fields of view per condition. Error bars indicate SEM at each time point. **(C)** Maximum intensity x-z projections of spinning disk confocal z-stacks showing FLS imaged with time. The initial and final images are the whole field of view, whilst the highlighted selection is a side projection of a 16×32 micron sub-region to track FLS behaviour during the timelapse. **(D)** FLS treated with R347 disassemble their bleached actin while with mAb14 treated FLS the bleached actin remains revealed by imaging at 25-35 min timepoints. Image panels as in (C). All scale bars = 10 μm.

The initial rate of assembly of actin during the growth phase of FLS (5 mins – 15 mins) was comparable across both R347 and mAbs 14, 2 and 6 conditions (Fig 5 B-C, S8A, Movie S1). However, FLS treated with lengthening antibodies displayed strongly reduced disassembly, as the bleached regions remained evident over the course of the imaging period (Fig 5C, S8A, Movies S1 and S2). The effect was even more apparent when imaging at steady state (25 – 35 mins), where the bleached actin disassembled with R347 and persisted with mAbs 14, 2 and 6. At this time point, the rate of freshly incorporated actin remained comparable to addition during the growth phase in the R347 treated FLS and could be slower with the lengthening antibodies (Fig 5C, S8B, Movies S1 and S2), likely due to the reduced availability of fresh monomers. Thus, a dynamic analysis of actin incorporation rates demonstrates that the lengthening effect of the anti-D-loop antibodies is due to the protection of FLS from disassembly.

### Additional cofilin rescues the altered length distribution of FLS

We prepared FLS with supplemental actin (providing extra antigen) and cofilin (as a D-loop binding disassembly protein) in the presence of the lengthening antibodies to determine whether either protein could suppress the lengthening effect or GFP-UtrCH displacement effect of mAb14. With the R347 control antibody, adding supplemental actin or cofilin did not displace GFP-UtrCH from FLS (Fig 6A) and increased actin caused lengthening of FLS (Fig 6A-B). In contrast, adding actin or cofilin had distinctly different effects on mAb14-treated FLS (Fig 6A,C). The addition of actin alone reduced the length of FLS and increased the degree of UtrCH-actin overlap, consistent with actin binding to mAb14 leading to a lowered effective antibody dose. Adding supplemental cofilin decreased FLS lengths with displacement of GFP-UtrCH remaining (Fig 6A,C). Supplemental actin in combination with cofilin had no additional effect on the length phenotype to cofilin alone, but as with the other instances of actin addition it restored approximately 50% of the fluorescent actin – GFP-UtrCH overlap (Fig 6D-E). Overall our data reveal that FLS are robustly lengthened when the open D-loop conformation of F-actin is stabilized or occupied by antibodies, which acts to prevent prevent disassembly of the actin bundle. This stabilization can be countered by supplementing the extracts with supplemental cofilin.

**Fig 6.**
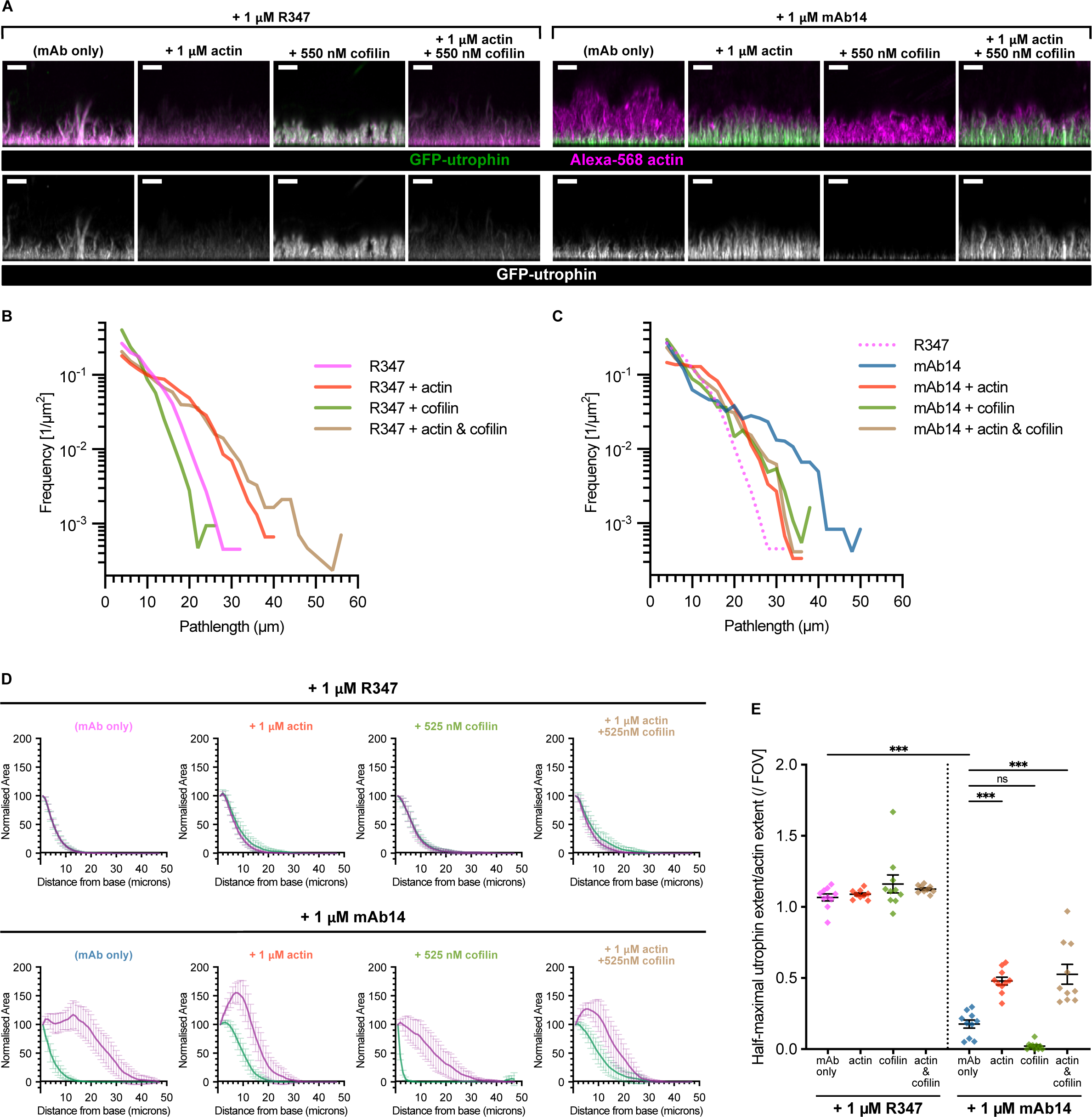
Supplementation of cofilin and actin rescues the effects of mAb14. **(A)** With mAb14, additional actin both shortens FLS and allows GFP-UtrCH binding. Additional cofilin shortens FLS and both cofilin and actin show synergistic effects. Maximum intensity x-z projections of spinning disk confocal z-stacks showing Alexa-568-actin (magenta) and GFP-UtrCH (green, single channel) in representative FLS regions preincubated with 1 μM NIP228 or mAb14 and supplemental actin, cofilin, or both as indicated. All scale bars = 10 μm. **(B)** Quantification reveals that with the control antibody, additional actin leds to longer FLS and cofilin alone slightly reduces FLS length. n(FLS) = 4417 (R347), 4541 (R347 + actin), 2135 (R347 + cofilin) & 4260 (R347 + actin + cofilin). **(C)** With mAb14 additional actin reduces the increase in length and increased cofilin both decreases length and restores the exponential distribution. n(FLS) = 2418 (mAb14), 2970 (mAb14 + actin), 1849 (mAb14 + cofilin) & 2417 (mAb14 + actin + cofilin). **(D)** Actin and UtrCH averaged fluorescence Z-slice profiles indicating the normalised relative fluorescence signal area at each Z slice for each channel averaged across each of N=9-10 imaging regions. Each slice is centered at the mean, error bars indicate 95% confidence intervals. **(E)** Quantification of GFP-UtrCH vs actin extent in each condition. Additional actin restores UtrCH staining to about half of the control mAb. Each datapoint represents an individual imaging region. N=9-10 fields of view per condition. Lines indicate mean ± SEM. Significance was assessed by ordinary one-way ANOVA with Sidak’s multiple comparison test: R347 only vs mAb14 only, mAb14 only vs mAb14 + actin & mAb14 only vs actin & cofilin, p < 0.001 [***], mAb14 only vs mAb14 + cofilin, p = 0.058 [ns]. Further comparisons can be found in Table S1.

## Discussion

We have identified a unique set of antibody reagents that provide insight into how actin conformational changes impact disassembly mechanisms of filopodia-like actin bundles. Structural transitions within the D-loop of actin filaments cause differential interactions of actin binding proteins, which in turn leads to the assembly of the large variety of architecturally distinct actin networks required for specific cellular functions. Actin filaments undergo conformational changes linked to the ATP hydrolysis that occurs on polymerization. In particular, the slow release of the inorganic phosphate group and the accompanying transition between open and closed D-loop state generates an age gradient along the filament linked to the nucleotide state, which can be used as signals for actin regulatory factor binding [21]. It is recognised that the proximal ends of parallel actin bundles, such as found in microvilli or stereocilia are enriched in ADP-actin interacting proteins [42]

The set of antibodies we identified disrupted the steady-state length distribution of FLS, which implies an alteration in the molecular mechanisms of length regulation. Actin-stabilizing compounds phalloidin and jasplakinolide can decouple D-loop conformational changes from the nucleotide state, trapping the filament in the closed or open state respectively [39]. Using phalloidin and jasplakinolide to stabilize these specific conformations revealed that the antibodies specifically recognise the open D-loop conformation, and displace the closed D-loop binding probes UtrCH and LifeAct. In doing so the antibodies prevented actin disassembly, which we could reveal by separating actin assembly and disassembly processes using photobleaching and timelapse imaging.

It is possible that the antibodies bind elsewhere on the bundle and prevent the probes binding by stabilsing the open D-loop conformation allosterically rather than by directly binding and occluding the D-loop. Molecular modeling and/or structural biology could be used to further elucidate the epitopes of the antibodies, initial molecular dynamics simulations have been inconclusive due to the size of the actin filament.

Addition of supplemental cofilin reduced the excess bundle length caused by the antibodies. Disassembly factors are particularly important as respondents to filament nucleotide status. In particular, cofilin and another disassembly factor coronin both bind at the D-loop and function in concert to induce disassembly, acting synergistically to enhance severing [43]. Cofilin preferentially binds ADP-actin (the aged closed conformation), inserting between the C-terminus and D-loop to alter the helical twist of the filament and ultimately induce severing. It requires the release of the inorganic phosphate group from the filament to function [20,27]. In contrast, coronin-1B recognises the ADP-Pi state around 50 times as strongly as it does the ADP state [21,37]. We have not distinguished whether cofilin directly competes with antibodies for D-loop binding or if it induces D-loop closure at high concentrations. Distinct from previous work on actin depolymerization studying small numbers of filaments, here we have been examining large fascin-bundled actin superstructures within a cytosolic-like environment of *Xenopus* egg extracts. Thus, this work accounts for the complex multiprotein interactions and effects of multiple actin filaments working together as a dynamic unit, and we have identified that the fundamental structural characteristics of the actin filament are the major functional contribution. In summary, our work implies that the transition to the closed state of the actin filament D-loop is a critical conformational switch that determines the lengths of filopodia-like F-actin bundles.

## Materials and Methods

This research has been regulated under the Animals (Scientific Procedures) Act 1986 Amendment Regulations 2012 after ethical review by the University of Cambridge Animal Welfare and Ethical Review Body and covered by Home Office Project Licence PP1038769 (licence holder: J.L. Gallop) and Home Office Personal licences held by H. Fisher, T. Jones-Green, and J. Mason.

### Antibodies, reagents and protein purification

The screen that isolated mAbs 2, 4, 6, 14 and 17 was described in (Jarsch et al., 2020) [29]. For each of these antibodies, as well as the R347 and NIP228 isotype controls, plasmids for expression of the HuIgG1 heavy chain and the IgG light chain were contransfected into CHO-transient cells and the IgGs purified from the cell culture medium via Protein A chromatography. DSHB antibodies A-E were obtained as hybridoma conditioned culture media supernatants from the Developmental Studies Hybridoma Bank, created by the NICHD of the NIH and maintained at The University of Iowa, Department of Biology, Iowa City, IA 52242. Antibody A is JLA20, deposited to the DSHB by Jim Jung-Ching Lin, Cold Spring Harbor Laboratory [44], Antibody B is CPTC-ACT Group 1, deposited by Clinical Proteomics Technologies for Cancer, National Cancer Institute, Antibody C is 224-236-1, deposited by Guenther Gerisch, Max-Planck Institute of Biochemistry [45], Antibody D is BB8/20.1 1, deposited by Belinda Bullard, University of York, and Antibody E is 8-7A5, deposited by Angel L. De Blas, University of Connecticut [46]. The mouse anti-pan actin antibody was from Cytoskeleton (#AAN02-S). The anti-human Alexa-647 conjugated secondary antibody (#109-605-008) and anti-mouse HRP conjugated secondary antibody (#715-035-150) were from Jackson ImmunoResearch. The anti-human HRP conjugated secondary antibody was from Southern Biotech (#9040-05). The anti-mouse IRDye CW800 conjugated secondary antibody was from LI-COR Biosciences (#926-32210) Atto-488 conjugated actin was from Hypermol (#8153-02). Atto-655-LifeAct was from Cambridge Research Biochemicals (#crb1101303h). Alexa-568 conjugated actin (#A12374), Alexa-488 coupled phalloidin (#A12379), unlabelled phalloidin (#P3457) and jasplakinolide (#J7473 ) were from Thermo Fisher Scientific. Purified rabbit skeletal muscle actin (#AKL99) and human recombinant cofilin (#CF01) were from Cytoskeleton. The GFP-UtrCH binding domain construct was originally a kind gift from Sarah Woolner, University of Manchester, and cloned into a pGEX vector for bacterial expression. This construct was transformed into Rosetta DE3 pLysS *E.coli* (Merck, #70956) and expression induced with IPTG overnight at 19°C. Cells were harvested by centrifugation and resuspended in 150 mM NaCl, 20 mM HEPES, 2 mM EDTA, 2 mM DTT and EDTA-free cOmplete protease inhibitor (Roche #11873580001), then lysed by probe sonication. The lysate underwent ultracentrifugation at 40,000 rpm for 45 mins in a 70Ti rotor (Beckman-Coulter), with the supernatant applied to 5 ml equilibrated Glutathione Sepharase 4B beads (Merck #17-0756) at 4 °C for 1 hour. Following five washes in 150 mM NaCl, 20 mM HEPES, 2 mM EDTA, 2 mM DTT, three washes in 300 mM NaCl, 20 mM HEPES, 2 mM EDTA, 2 mM DTT and 2 washes in 150 mM NaCl, 20 mM HEPES, 2 mM EDTA, 2 mM DTT, 25mM CaCl_2_, the beads were resuspended in 2 ml of the latter buffer with 100 units Thrombin (GE Healthcare #27-0846) and cleaved at 4 °C overnight. The beads and supernatant were separate via a gravity flow column, and the purified GFP-UtrCH separated from the thrombin by passing over a HiTrap Q FF column (#GE17-5156) on an ÄKTA FPLC system (both Cytiva) in a buffer containing 150 mM NaCl, 20 mM HEPES, 2 mM EDTA, 2 mM DTT. The elutions were pooled and concentrated in an Amicon Ultra 10 kDa MWCO spin concentrator (Merck #UFC801024), then diluted in 50% glycerol and stored at -20 °C.

### FLS assays

*Xenopus laevis* egg extracts, supported lipid bilayers composed of 60% phosphatidylcholine (#840053C), 30 % phosphatidylserine (#840032C) and 10% PI(4,5)P_2_ (#840046X, all Avanti Polar Lipids), and FLS assays were performed as previously described [47]. For all experiments, a final 1:6 dilution of 25 mg/ml extracts was used in a 50 µl volume reaction mix also containing 2 mM DTT, energy mix (50 mM phosphocreatine, 20 mM Mg-ATP) and made up in extract buffer (XB) (100 mM KCl, 100 nM CaCl_2_, 1 mM MgCl_2_, 10 mM K-HEPES pH 7.4, 50 mM sucrose). When required (and as indicated in Figs), GFP-UtrCH and 655-LifeAct were added to the reaction mix at 250 nM, fluorescent actins were added at 210 nM, supplemental unlabelled actin was added at 1 µM, and cofilin was added at 550 nM. Antibodies were exchanged into a buffer containing 100 mM KCl, 1 mM MgCl_2_, and 10 mM K-HEPES pH 7.4 via a 10 kDa MWCO spin concentrator, and added at the indicated concentrations to the FLS mix alongside an equal volume of double-sucrose XB (XB as above containing 100 mM sucrose rather than 50 mM). DSHB antibodies were buffer exchanged as above and 8 µl added to the reaction mix, alongside an equal volume of double-sucrose XB. Phalloidin and jasplakinolide were prepared as 2.5 mM stocks in methanol, and added to the reaction mix at the indicated concentrations. The vehicle control was methanol added at a 1:500 final dilution, corresponding to the volume required for the highest phalloidin and jasplakinolide concentration. Each intervention was allowed to pre-incubate for 10 mins before being added to the supported lipid bilayer to initiate the assay. Steady state snapshots were imaged over the time period 25-35 mins after assay initiation. Timelapse experiments were imaged over the time periods 5-15 mins and 25-35 mins after initiation. For FLS immunolabelling experiments, assay mixes in XB containing frog egg extracts, DTT, energy mix, Atto-488 actin were prepared and initiated as above. At T=20 minutes, 0.5 µl phalloidin from a 10 mg/ml methanol stock was added for a further 5 minutes. The structures were gently washed twice with 50 µl XB containing 1x energy mix and 2mM DTT, then fixed in 50 µl 4% formaldehyde (#28908, Thermo Fisher Scientific) in XB for 75 mins. 50 µl 50 µM glycine in PBS was added for 15 mins to quench the fixative, then the structures were blocked for 60 mins in 10% goat serum (Merck #G6767) in PBS. Primary antibodies were prepared at a 1:100 dilution (corresponding to final concentrations between 40-65 nM) in 1% goat serum in PBS, and 50 µl added to each well, then incubated at 4 °C overnight. Assays were carefully washed in 1% goat serum-PBS three times for five minutes. The anti-human Alexa-647 secondary antibody was prepared at a 1:200 dilution in 1% goat serum-PBS, added, and incubated for 1 hour. Assays were carefully washed in 1% goat serum-PBS three times for five minutes and immediately imaged.

### FLS Imaging

FLS were imaged at room temperature on a combined spinning disk confocal/TIRF system provided by CAIRN research equipped with an Eclipse-Ti inverted microscope (Nikon), X-light Nipkow spinning disk unit (CREST), Spectra X LED illuminator (Lumencor) and NanoScanZ 250 µm piezo stage (Prior) via a 100x 1.40 NA Plan Apo VC oil objective (Nikon) on an Evolve Delta EMCCD camera (Photometrics) in 16-bit depth using Metamorph software. Alexa/Atto 488, 568 and 647/655 samples were visualised using 470/40, 560/25 and 628/50 excitation and 525/50, 585/50 and 700/75 emission filters respectively. FLS were imaged at one micron Z-stack intervals, taking 30-60 micron stacks, with the exception of the immunolabelling and live imaging experiments, which were both imaged at 0.5 micron intervals. For bleaching experiments, an iLas2 illuminator (Roper scientific) via a 100x 1.49 NA oil immersion objective was used to bleach the actin channel in an entire field of view at the level of the supported lipid bilayer via the TIRF laser (100% power, 3 repetitions). Images were taken via the standard confocal set up used in all other experiments. At the correct timepoint after initiation, a reference 60 micron z stack at 0.5 micron intervals was taken before bleaching, followed by bleaching at the base slice, as described above. An immediate time-lapse was initiated, capturing a 26 micron z stack at 0.5 micron intervals every 20s for 10 mins. A second 60 micron z stack at 0.5 micron intervals was then taken at the conclusion of the timelapse. For each assay, a region of interest was captured between 5-15 minutes after assay initiation, then a second region of interest was used to assay the 25-35 minute timepoint.

### Cell culture and immunolabelling

HEK293 cells (ATCC) were grown as adherent cultures at 37 °C in a 5% CO_2_ atmosphere. They were maintained in Dulbecco’s modified Eagle’s medium (Merck #D6546) supplemented with 10% fetal bovine serum (Merck #F9664) and 1X penicillin-streptomycin-glutamine (Thermo Scientific #10378016), and passaged twice weekly. They were passaged onto glass coverslips, and allowed to grow to approximately 50% confluency before processing. Coverslips were fixed in 4% formaldehyde for 10 minutes, then washed three times in PBS. They were permebilised and blocked in a mix containing 0.1% saponin, 10% goat serum and 50 mM NH_4_Cl in PBS for 45 minutes. Primary antibody mixes containing 4 µg/ml (approximately 25 nM) of each relevant antibody were prepared in 1% goat serum-PBS, and incubated on the coverslips overnight at 4 °C. They were then washed three times at 4 °C in 1% goat serum-PBS and incubated with a mix containing Alexa-488 phalloidin (1:250) and the anti-human alexa-647 secondary antibody (1:500) in PBS for 45 mins at 4 °C. They were then washed four times at 4 °C in 1% goat serum-PBS, and mounted using Hydromount (National Diagnostics #HS-106). Images were acquired on an Airyscan Zeiss LSM 880 via a Plan-Apochromat 63x NA 1.4 objective using the Airyscan Fast imaging mode as a confocal z-stack taken at 0.2 µm intervals. Images were processed using Airyscan processing, strength 6, and are presented as maximal z-projections.

### *Xenopus laevis* embryo injection and cytokinesis assay

Freshly laid *X. laevis* eggs were fertilized and dejellied using 2% cysteine in MMR, then transferred into 4% Ficoll-0.1% MMR for microinjection. Once embryos reached the two cell stage, one cell in each embryo was injected with 10 nl of 9 µM Ab14 or R347 using a Parker Picospritzer III microinjector. Embryos were monitored and on a Zeiss Stemi 305 stereo microscope, with images captured at each division stage. Embryos for each antibody at the 4 and 8 cell stage were scored for defective cytokinesis by two independent researchers, and the mean of those values reported.

### SDS-PAGE and Western blotting

Samples were prepared in 4x Laemmli sample buffer (final 1x composition: 2% SDS, 50mM Tris pH 6.8, 10% glycerol, 5% beta-mercaptoethanol) and run on 4-20% mini-PROTEAN TGX precast polyacrylamide gels (Bio-Rad #4561096) in Tris-glycine-SDS running buffer, using the PageRuler Plus Prestained Protein Ladder (Thermo Fisher #26619) as a molecular weight marker. For coomassie staining, InstantBlue stain (abcam #ab119211) was added to gels overnight, then destained in ddH_2_O prior to scanning. For the anti-actin Western blot, gels were transferred onto nitrocellulose membrane using the iblot 2 dry blotting system (Thermo Fisher #IB23002). Membranes were blocked in 10% milk (Merck #70166) in TBST, then incubated with the anti pan actin primary antibody overnight at 4 °C (dilution 1:500 in 1% milk-TBST). Following three washes in 1% milk-TBST, the membranes were incubated in with the anti-mouse CW800 secondary antibody (1:5000 in 1% milk-TBST), followed by three further washes in 1% milk-TBST before scanning. Both coomassie gels and Western blots were scanned on an Odyssey Sa reader (Li-COR biosciences). Densitometry analysis was performed on scanned coomassie gel images using FIJI (ImageJ) software [48]. The intensity of each band representing actin or the antibody heavy chain in the pellet and supernatant for each treatment was measured, as well as two strips above and below these bands to measure the background. The average of these strips was taken and subtracted from each individual band measurement. For each condition the percentage of protein in the pellet was calculated from these background subtracted measurements as: Pellet Intensity / (Pellet Intensity + Supernatant Intensity) x 100.

### Immunoprecipitation assays

Antibodies were pre-cleared by centrifugation at 5000 xg for 5 mins at 4 °C. For each condition, 24 µg of relevant antibody was coupled to 100 µl protein A dynabeads in PBS for 2 hours at 4 °C on a rotator, after which beads were washed twice in 1ml 0.02% Tween20-PBS (PBST), once in 1 ml PBST + 250 mM NaCl, and twice in 1 ml XB. Meanwhile, for each condition, 100 µl *X. laevis* egg extracts were preincubated with either 20 µM Latrunculin-B (Merck, #428020), or the equivalent volume of DMSO (vehicle control) for 10 mins on ice. The washed antibody-coupled beads were incubated with the relevant pretreated frog egg extracts for 4 hrs at 4 °C on a rotator. They were then washed in 1 ml XB followed by 3 x 1 ml washes in PBST. The beads were then resuspended in 30 µl Laemmli sample buffer and heated at 95 °C in a heat block for 5 mins. The supernatant was separated from the beads and run on gels for coomassie staining and Western blotting.

### Sedimentation assays

Rabbit skeletal muscle actin was resuspended to 1 mg/ml in general actin buffer (5 mM Tris-HCl pH 8.0, 0.2 mM CaCl_2_, 0.2 mM ATP, 0.5 mM DTT). Antibodies were diluted to 0.6 µg/ml in PBS. Both underwent a clearing spin at 150,000 xg for 1 hr at 4 °C. For each assay, 52.5 µl of cushion buffer (2.5 µl 10x actin polymerization buffer (Cytoskeleton #BSA02), 7 µl glycerol, 2 µl ddH_2_O, 1 µl 10x PBS, 40 µl general actin buffer) was prepared and added to the bottom of the ultracentrifuge tube (2.5 µl 10x polymerization buffer [500 mM KCl, 20 mM MgCl_2_, 10 mM ATP]. On top of this the reaction mix was carefully overlaid as follows: 40 µl cleared actin, 2.5 µl 10x polymerization buffer, 10 µl ( µg) cleared antibody. Assays were incubated at 4 °C for 1 hr, then centrifuged at either 150,000 xg for 90 mins (for the F-actin assay), or 14,000 xg for 60 mins (for the bundled filament assay). The supernatant was carefully removed and 30 µl laemmli sample buffer added. The pellet was then resuspended in 105 µl PBS and 30 µl laemmli sample buffer added. Both supernatant and pellet samples were heated in a 95 °C heatblock for 5 mins, then 10 µl of each sample run an an SDS-PAGE gel for coomassie staining.

### Actin ELISA assays

Rabbit skeletal muscle actin was resuspended to 1 mg/ml (21 µM) in 5mM Tris-HCl ph 8.0 + 0.2 mM CaCl_2_. Equal volumes were split to prepare F-actin or G-actin. G-actin was prepared by adding 0.2mM ATP and leaving on ice for 30 minutes, before adding a 1:10 volume of 5mM Tris-HCl ph 8.0 + 0.2 mM CaCl_2_ and leaving on ice for a further 30 mins. F-actin was prepared by adding a 1:10 volume of 10x actin polymerization buffer and leaving at room temperature for 1 hour. For jasplakinolide and phalloidin treatments, F-actin was prepared as above and left at room temperature overnight, before jasplakinolide or phalloidin was added to a final concentration of 40 µM. A dilution series of actin concentrations was prepared with G-actin diluted in 5mM Tris-HCl ph 8.0 + 0.2 mM CaCl_2_ + 0.2 mM ATP, and F-actin (and F-actin treated with jasplakinolide or phallodin) diluted in 1x actin polymerization buffer diluted in 5mM Tris-HCl ph 8.0 + 0.2 mM CaCl_2_. 50 µl volumes of each dilution were added to wells of a MaxiSorp 96-well Immuno plate (Thermo Scientific #437111) in triplicate and incubated overnight, with F-actin plates at room temperature and G-actin plates at 4 °C. The plates were then washed in 200 µl PBS-0.1% Tween-20 three times, then blocked in 200 µl 3% BSA in PBS for 1 hour. Antibodies were prepared in 1% BSA-PBS. R347 and mAbs 2, 6, 14 and 17 were used at 10 nM final concentration, the pan actin antibody was used at 1:1000. Wells were washed in 200 µl PBS-0.1% Tween-20 three times, then 100 µl of the primary antibody mixes added and incubated for 1 hour. Anti-human or anti-mouse HRP antibodies were prepared in 1% BSA-PBS at 1:10,000 dilution. Wells were washed in 20-0 µl PBS-0.1% Tween-20 three times, then 100 µl secondary antibodies mixes added and incubated for 1 hour, followed by three final 200 µl PBS-0.1% Tween-20 washes. HRP was detected using the QuantaBlu Fluorogenic peroxidase kit (Thermo Scientific #15169). The substrate and peroxidase solutions were mixed according to manufacturers instructions, and 50 µl added to wells for 30 mins, followed by the addition of 50 µl QuantaBlu stop solution. Plates were read on a Hidex Sense using 320/40 and 444/12 excitation and emission filters, or on a Perking Elmer Multilabel EnVision microplate reader. Response curves were calculated and plotted using the [Agonist] vs. response –Variable slope analysis in Graphpad Prism software (version 10.1.0).

### Quantification and statistical analysis

#### FLS lengths

FLS length data was measured using our custom FIJI (Image J) plugin FLSACE, described in [5], which maps FLS through Z-stacks. Lengths reported are the path-length of the segmented structure measured from the base slice. Structures are defined as FLS when they have a length of at least 3 µm, a base area of at most 20 µm^2^, and a base circularity of at least half that of a circle.

#### Actin vs actin probe binding

To calculate UtrCH and LifeAct displacement, the total area of fluorescence from the base slice upwards was measured in each z-slice in both channels of the confocal stack. In the actin channel, the actin extent was defined as the first z-slice with an actin area of zero. The UtrCH or LifeAct extent was defined as the first slice where the UtrCH area is less than or equal to half the actin area in that slice. This ratio of UtrCH or LifeAct extent / actin extent was calculated for each field of view and plotted as the half-maximal UtrCH extent / actin extent. The area of fluorescence measurements are normalised and plotted as the averaged fluorescence Z-slice profiles. For every Z-slice in a field of view, the fluorescence area measurement was normalised relative to the area of that channel in the first slice above the base slice. The graphs plot the mean normalised area at each z-slice for both channels.

#### Time lapses

To plot the timelapse mean length profiles, the lengths of individual FLS were tracked over time in FLSACE, with T=0 for each individual FLS set as its first detection by FLSACE. To be included, FLS had to be detected for a minimum of 6 consecutive frames (2 mins). The graph plots the mean length of all detected FLS at each time point relative to this first appearance. All graphs were prepared and all statistical analysis performed in Graphpad Prism software (version 10.1.0), as described in Fig legends.

---

**Movie S1. mAb14 protects FLS against depolymerization**

Maximal intensity x-z projection of spinning disk confocal time-lapse movie of FLS treated with either R347 or mAb14 immediately following photobleaching. The top panel illustrates examples bleached at T=5 mins of FLS growth, whilst the bottom panel shows examples bleached at T=5 mins of FLS growth. For each, images were captured every 20s immediately post photobleaching for a 10 minute period, and are replayed at 10 frames per second. Initial growth rates between control and mAb14 treated FLS are similar at the 5-15 minute timepoint. At steady state, incorporation of fresh actin is slower following mAb14 pre-incubation, however FLS are longer as the bleached actin has been protected from loss by depolymerization.

**Movie S2. mAb2 & mAb6 protect FLS against depolymerization**

Maximal intensity x-z projection of spinning disk confocal time-lapse movie of FLS treated with either R347, mAb2 or mAb6 immediately following photobleaching. The top panel illustrates examples bleached at T=5 mins of FLS growth, whilst the bottom panel shows examples bleached at T=5 mins of FLS growth. For each, images were captured every 20 s immediately post photobleaching for a 10 minute period, and are replayed at 10 frames per second. Initial growth rates between control and mAb2 and 6 treated FLS are similar at the 5-15 minute timepoint. At steady state, incorporation of fresh actin is slower following mAb2 pre-incubation, whilst at both timepoints mAb2 and mAb6 protect the bleached action from loss by depolymerization.

**Fig S1.**
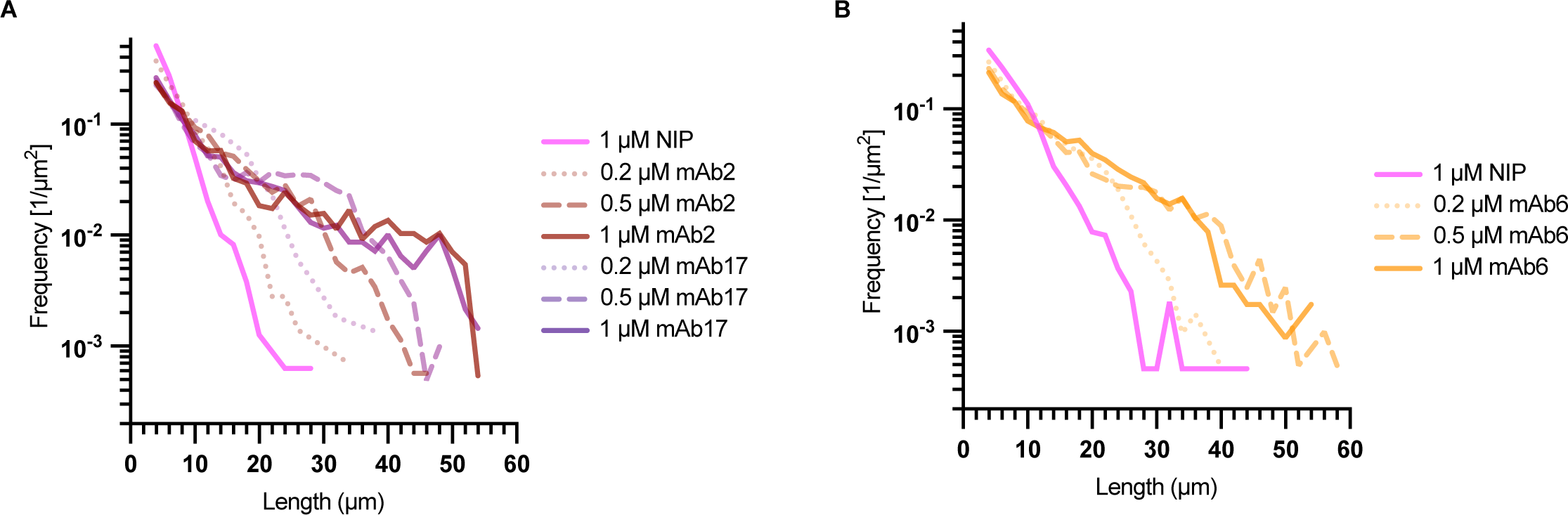
FLS lengthening mAbs shift the length distribution of FLS in a dose-responsive manner. Log FLS length histograms showing the dose responsive shift in FLS length distribution for **(A)** mAb2 and mAb17 and **(B)** mAb6, in both cases compated to 1 µM NIP228 treatment. **(A)** n(FLS) = 1430 (0.2 µM mAb2), 1754 (0.5 µM mAb2), 1840 (1 µM mAb2), 2189 (0.2 µM mAb17), 2021 (0.5 µM mAb17), 1385 (1 µM mAb17), & 1579 (1 µM NIP), **(B)** n(FLS) = 2096 (0.2 µM mAb6), 2026 (0.5 µM mAb6), 1150 (1 µM mAb6), & 2176 (1 µM NIP).

**Fig S2.**
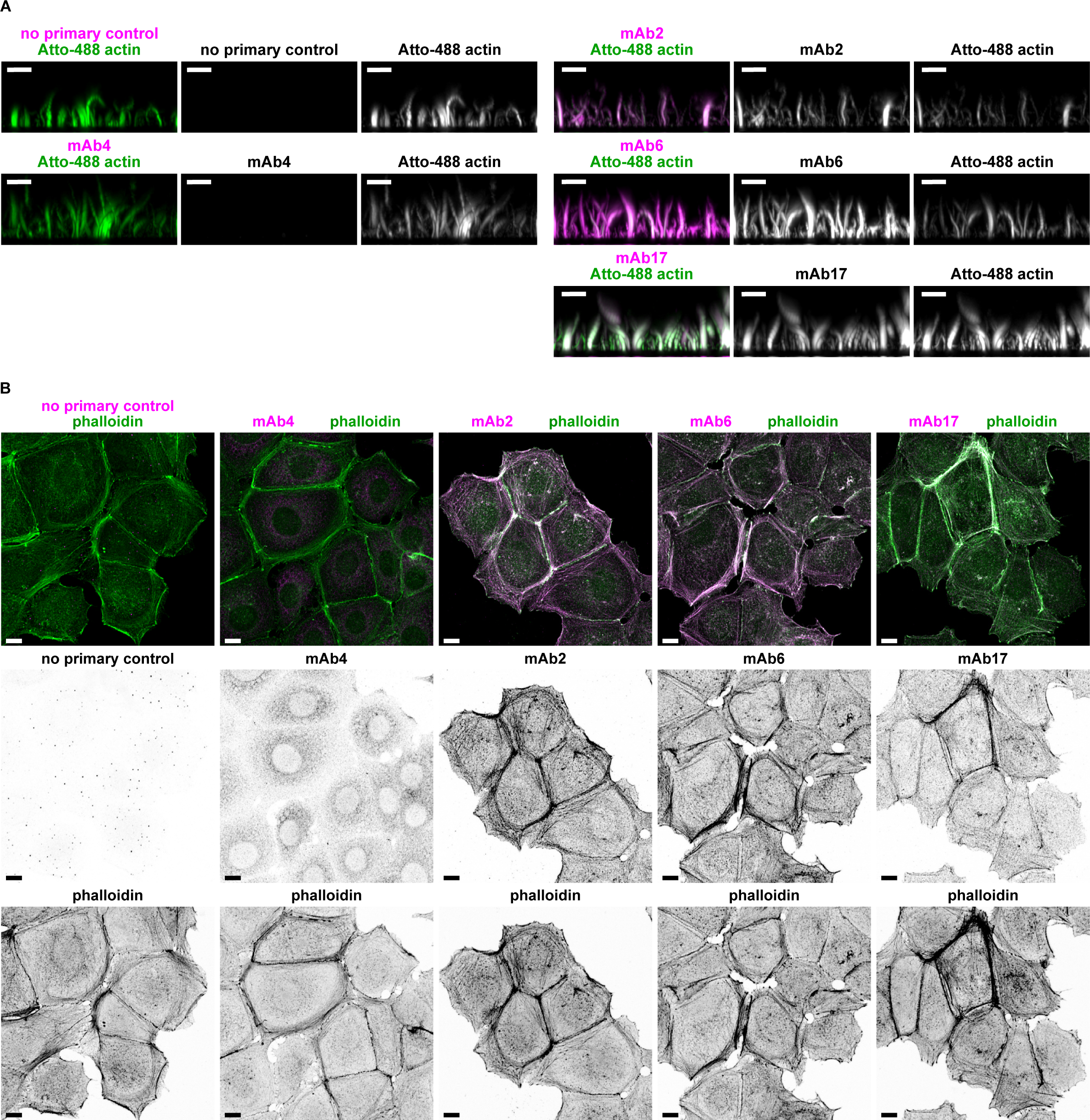
The FLS lengthening mAb group immunostains F-actin. **(A)** Maximal intensity x-z projection of spinning disk confocal z-stack of Atto-488-actin (green) in representative untreated FLS regions immunolabelled for mAb2, 6 or 17, mAb4, or the secondary antibody only (magenta). The FLS lengthening mAbs (2, 6 and 17) all label along the length of the FLS bundle, while anti-SNX9 antibody mAb4 (identified from the same phenotypic screen) does not. The lack of signal in the secondary control channel confirms the specificity of mAb labelling. **(B)** Maximal intensity x-z projection of laser scanning confocal images HEK293T cells immunolabelled for mAb2, 6 or 17, mAb4, or the secondary antibody only (magenta) and Alexa-488--phalloidin (green). The FLS lengthening mAbs (2, 6 and 17) all label along the F-actin, while anti-SNX9 antibody mAb4 labels discrete cytosplasmic puncta. The lack of signal in the secondary control channel confirms the specificity of mAb labelling. All scale bars = 10 μm.

**Fig S3.**
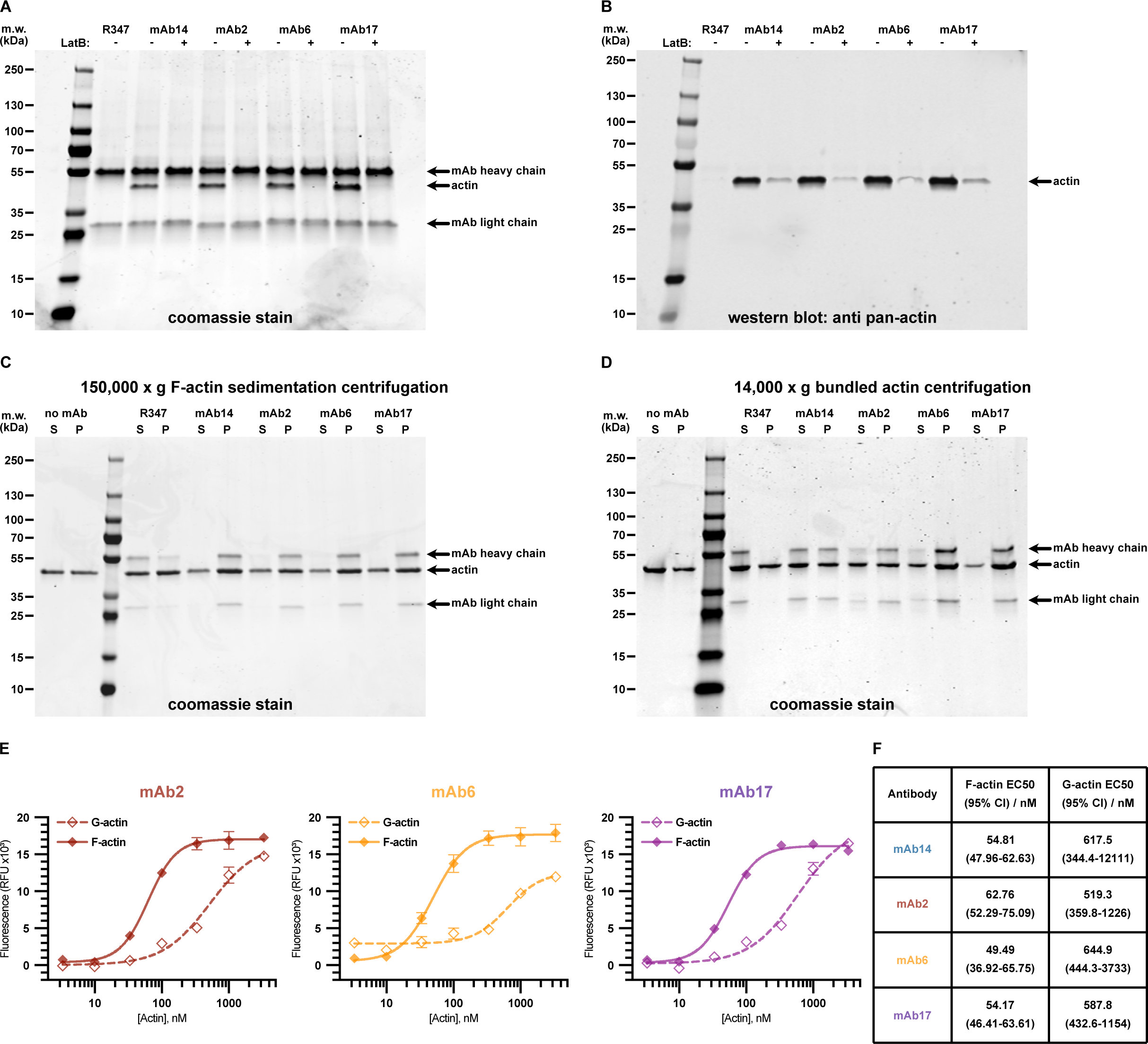
FLS lengthening mAbs bind to F-actin with greater affinity than G-actin. **(A-D)** Full gels and Western blots of the panels shown in Fig 2A and 2B. **(A)** is Fig 2Ai, **(B)** is Fig 2Aii, **(C)** is Fig 2Bi, and **(D)** is Fig 2Bii. **(E)** Variable slope [Agonist]:Response fitted binding curves for mAb2, 6 and 17 binding to F-actin vs G-actin. Symbols and error bars indicate the mean and SEM of each measured concentration (each data point was taken in triplicate). **(F)** Table outlining EC50 (and 95% confidence interval) values calculated from Variable slope [Agonist]:Response fitted binding curves for mAb14 (Fig 2E) and mAb2, 6 and 17 (E) antibody binding to F or G-actin. In each case the mAb binds to F-actin with an approximately 10-fold enhanced affinity than it binds G-actin.

**Fig S4.**
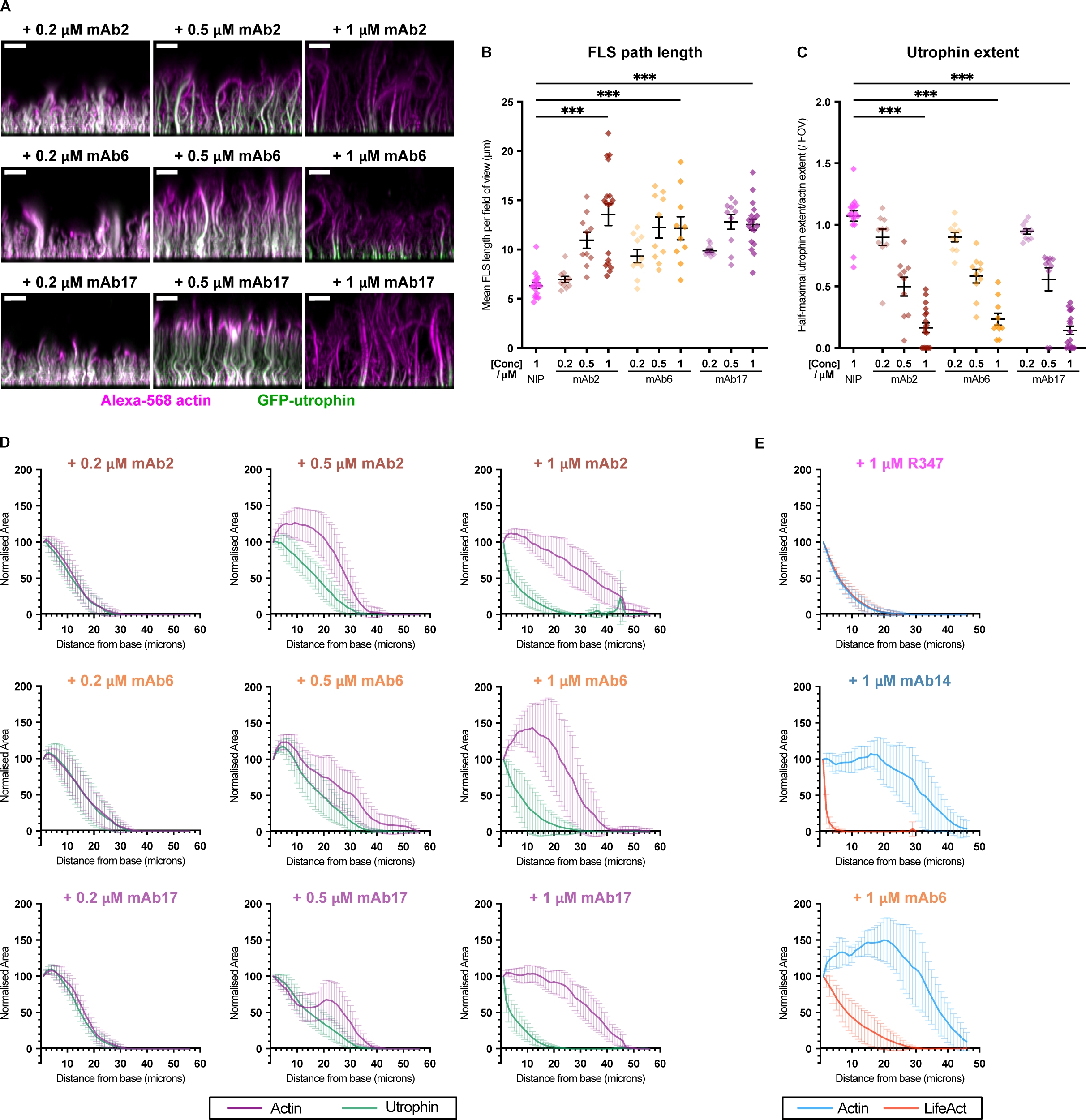
FLS lengthening mAbs displace the actin binding probe GFP-UtrCH domain from FLS actin bundles in a dose responsive manner. **(A)** Maximal intensity x-z projection of spinning disk confocal z-stack of Alexa-568-actin (magenta) and GFP-UtrCH (green, single channel) in representative FLS regions preincubated with 0.2, 0.5 or 1 μM mAb2, 6 and 17. All scale bars = 10 μm. **(B)** Quantification of average FLS length / FOV in each condition. Lengths significantly increase on the addition of FLS lengthening antibodies relative to 1 μM NIP228 control mAb, and do so in a dose responsive manner up to 0.5 μM of mAb6 and 17 and 1 μM of mAb2. Each datapoint represents an individual imaging region. N=10-18 imaging FOVs per condition. Lines indicate mean ± SEM. Significance was assessed by ordinary one-way ANOVA with Sidak’s multiple comparison test: NIP228 vs 1 µM mAb2, 1µM mAb6 & 1µM mAb17, all p < 0.001 [***]. Further comparisons can be found in Table S1. **(C)** Quantification of z-extent of GFP-UtrCH in each condition. GFP-UtrCH is displaced on the addition of each of mAb2, 6 and 17 in a dose responsive manner. Each datapoint represents an individual imaging region. N=10-17 fields of view per condition. Lines indicate mean ± SEM. Significance was assessed by ordinary one-way ANOVA with Sidak’s multiple comparison test: NIP228 vs 1 µM mAb2, 1µM mAb6 & 1µM mAb17, all p < 0.001 [***]. Further comparisons can be found in Table S1. **(D)** Actin and GFP-UtrCH averaged fluorescence Z-slice profiles indicating the normalised relative fluorescence signal area at each Z slice for each channel averaged across each of N=8-18 fields of view for 0.2, 0.5 or 1 μM of NIP228, mAb14, mAb2, mAb6 and mAb17. Each slice is centered at the mean, error bars indicate 95% confidence intervals. The two channels closely align on R347 treatment, whereas the GFP-UtrCH signal and distance from the base slice of the actin curves respond in a dose responsive manner for each of the mAbs. **(E)** Actin and LifeAct averaged fluorescence Z-slice profiles indicating the normalised relative fluorescence signal area at each Z slice for each channel averaged across each of N=6-8 fields of view for 1 μM of R347, mAb14, and mAb6. Each slice is centered at the mean, error bars indicate 95% confidence intervals. The two channels closely align on R347 treatment, whereas the LifeAct signal is substantially shorter on mAb14 or 6 treatment, where the actin is detected at much greater distance from the base. Representative images and quantification are shown in Fig 3 E and F.

**Fig S5.**
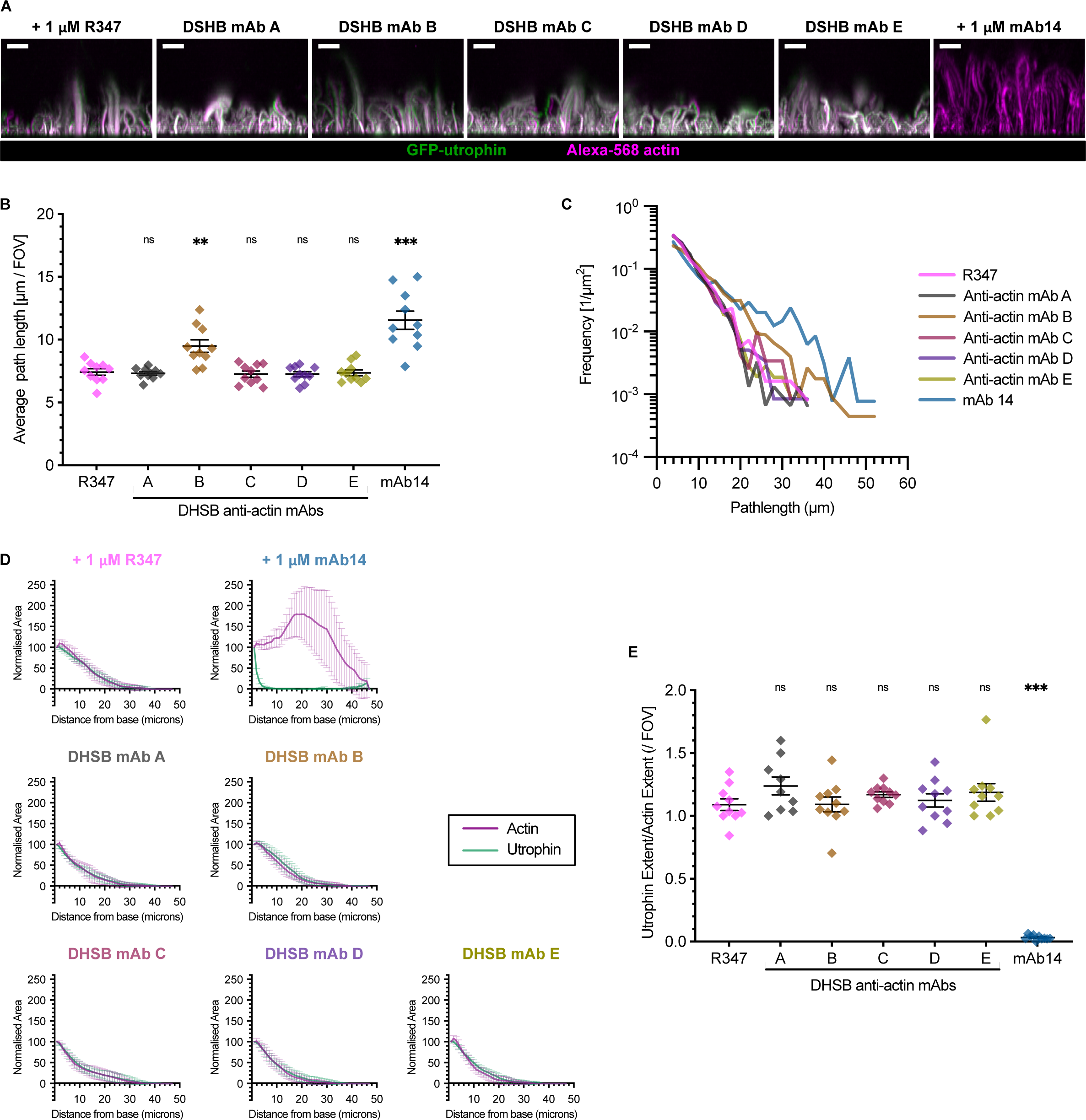
FLS lengthening is not a common property of anti-actin antibodies. **(A)** Maximal intensity x-z projection of spinning disk confocal z-stacks showing GFP-UtrCH and actin from anti-actin antibodies from the Developmental Studies Hybridoma Bank. **(B, C)** The lengths of most remain similar with a slight increase with an antibody B which was raised against a peptide comprising residues 29-39 common to all actin isoforms. **(D, E)** GFP-UtrCH is not displaced by most anti-actin antibodies. N=9-10 imaging regions per condition. Significance was assessed by ordinary one-way ANOVA with Sidak’s multiple comparison test and is reported in Table S1.

**Fig S6.**
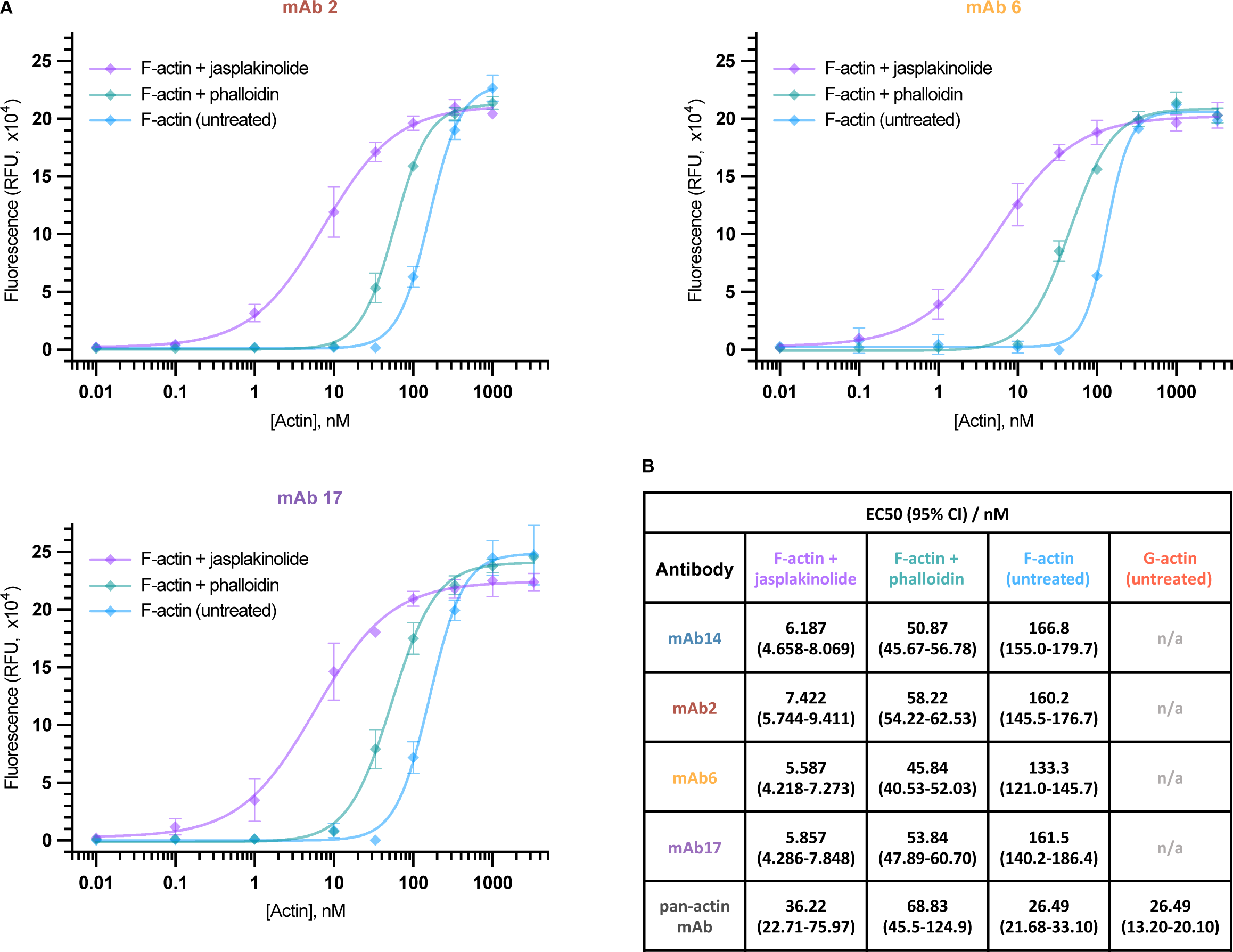
FLS lengthening mAbs specifically recognise F-actin the open conformation of the actin D-Loop. For each of mAb2, 6, 14 and 17 the mAbs bound with an approximately 10 fold greater affinity to F-actin locked in the open conformation via jasplakinolide compared to being locked in the closed conformation via phalloidin treatment. The pan-actin mAb bound with equal EC50 values to untreated F and G-actin, and with decreased affinity to either jasplakinolide or phalloidin stabilized filaments. **(A)** Variable slope [Agonist]:Response fitted binding curves for mAb2, 6 and 17 binding to untreated aged F-actin, and aged F-actin treated with jasplakinolide or phalloidin. Symbols and error bars indicate the mean and SEM of each measured concentration (each data point was taken in triplicate). **(B)** Table outlining EC50 (and 95% confidence interval) values calculated from variable slope [Agonist]:Response fitted binding curves for mAb14 (Fig 4B) and mAb2, 6 and 17 (A) antibody binding to untreated aged F-actin, and aged F-actin treated with jasplakinolide or phalloidin, and the pan-actin mAb binding to the same three treatments plus untreated G-actin.

**Fig S7.**
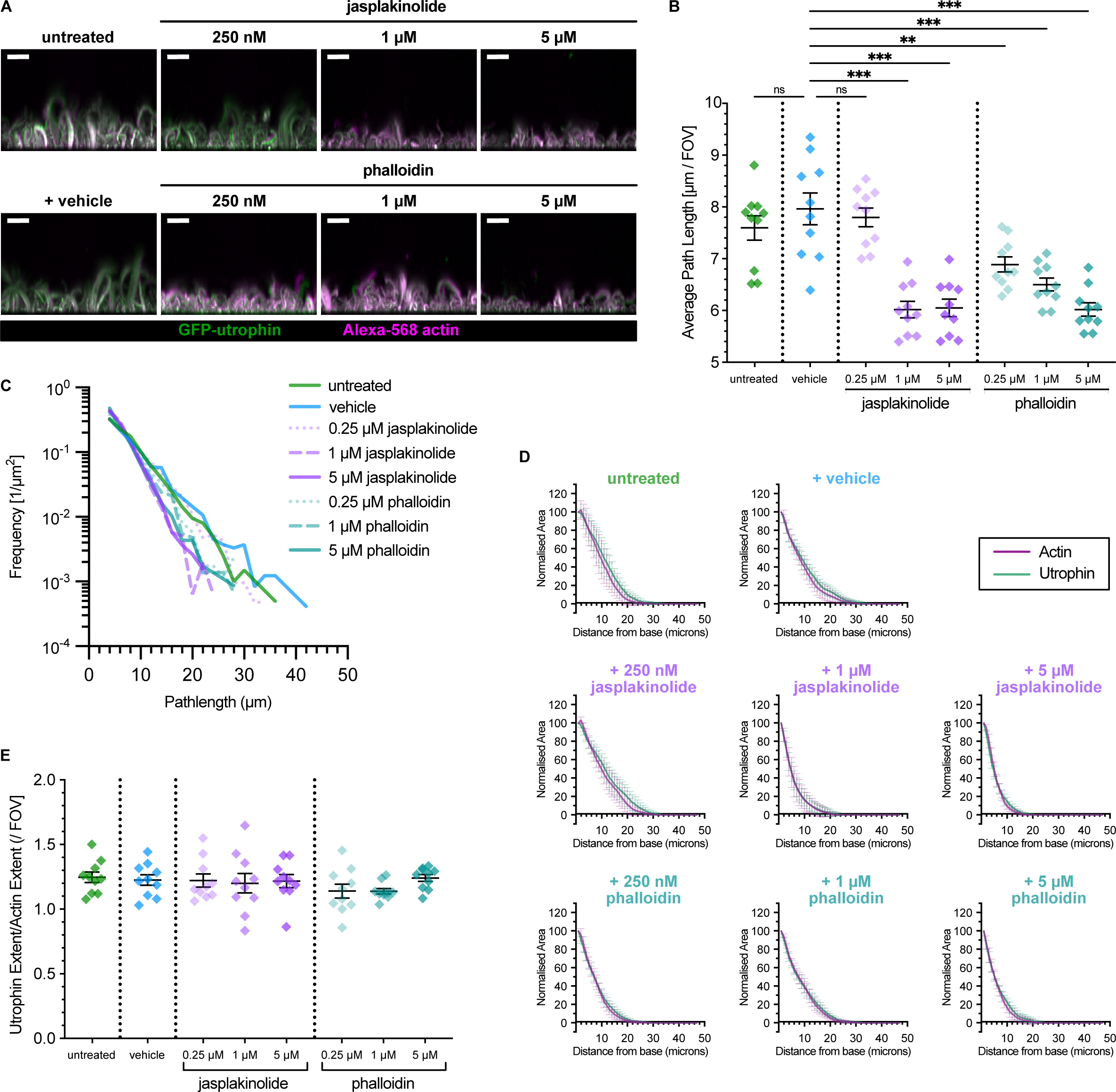
Jasplakinolide and phalloidin reduce FLS lengths and have no effect on GFP-UtrCH binding. **(A)** Maximal intensity x-z projection of spinning disk confocal z-stacks showing GFP-UtrCH and actin labelling FLS in the presence of jasplakinolide and phalloidin. **(B)** The mean length of FLS is reduced in a dose-responsive manner. **(C)** The length distribution of FLS is similar between conditions. **(D, E)** There is no difference in fluorescence extent between actin and GFP-UtrCH labelling of FLS. N=10 imaging regions per condition. Significance was assessed by ordinary one-way ANOVA with Sidak’s multiple comparison test and is reported in Table S1.

**Fig S8.**
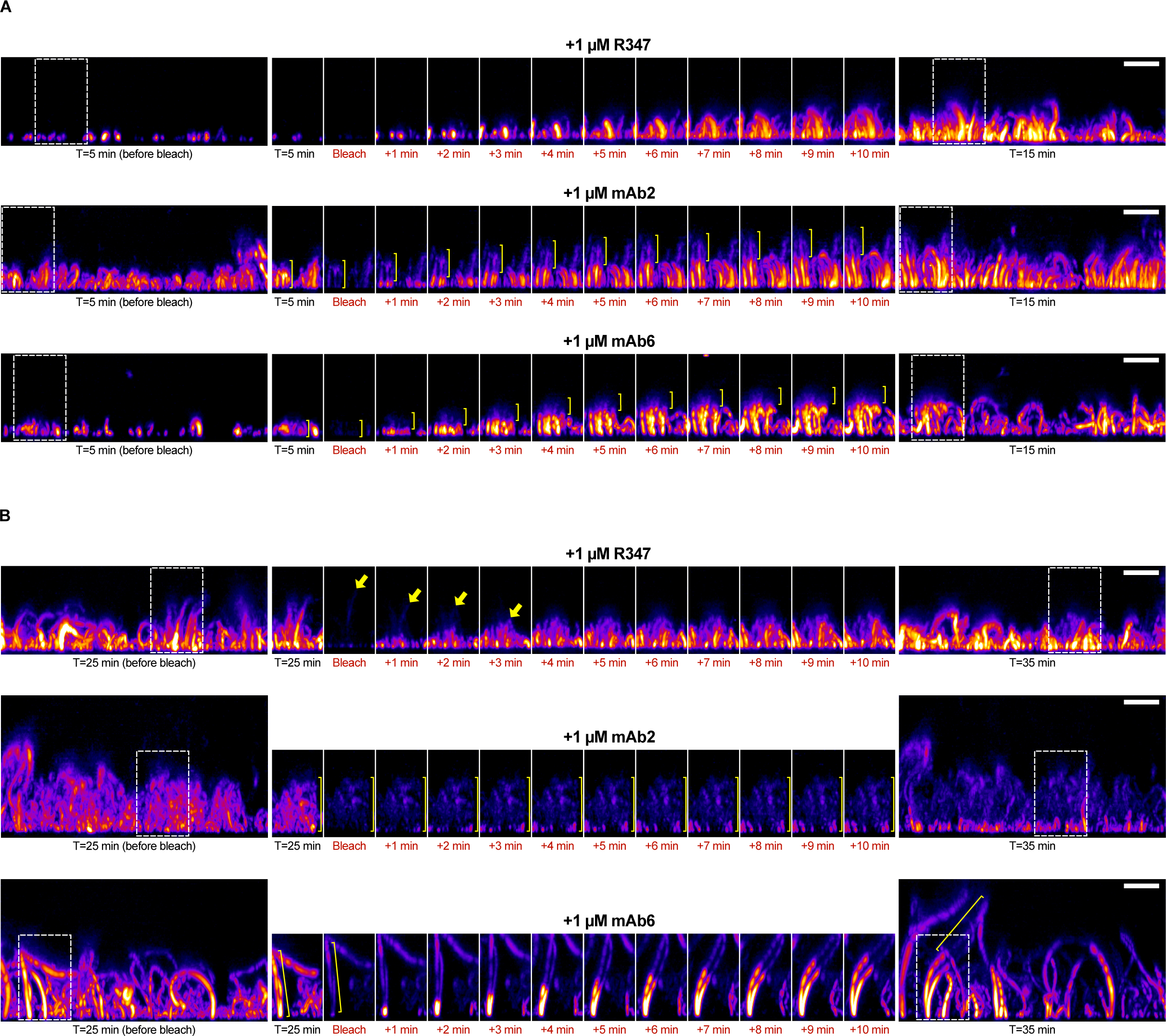
FLS lengthening mAbs protect actin bundles from disassembly. **(A-B)** Maximal intensity x-z projection of spinning disk confocal z-stacks with labelled actin in the presence of either R347, mAb2 or mAb6 imaged before photobleaching, and every minute for 10 minutes after photobleaching at the **(A)** T=5-15min, and **(B)** T=25-35min timepoints. In each panel, the initial and final images illustrate the full field of view, and the highlighted selection is a side projection of a 16 x 32 micron sub-region to track FLS behaviour during the timelapse. In the 5-15min timelapse the incorporation rates are similar, however protected regions of bleached actin remain with both mAb2 and mAb6 (marked). At the 25-35 min timepoints the incorporation rate can vary by region, and is often slower with antibody treatment, however the bleached regions (highlighted) remain and do not depolymerize, in contrast to the R347 treatment, where the bleached actin is lost over the time-course (yellow arrows). All scale bars = 10 μm.

**Table S1.**
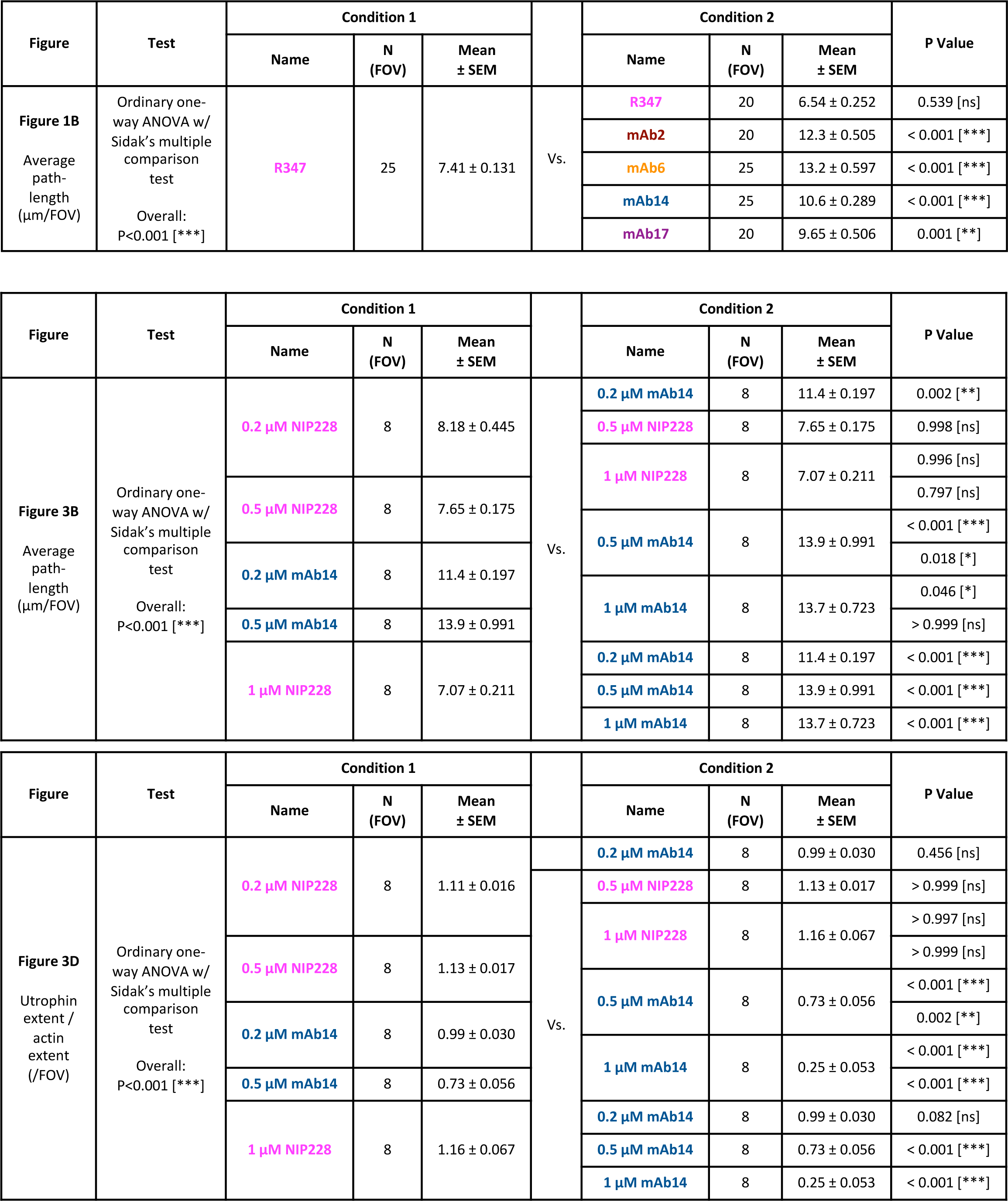
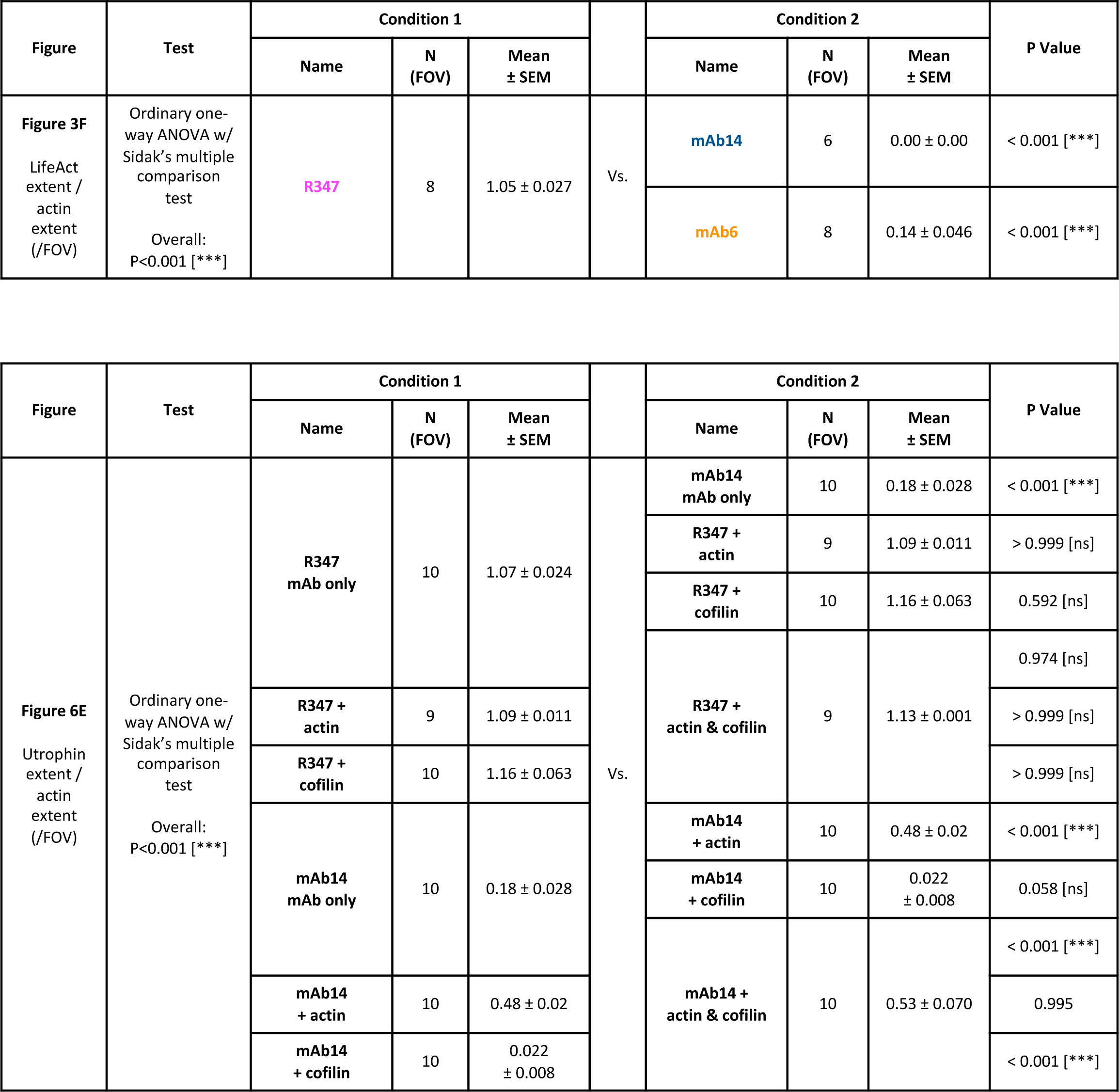
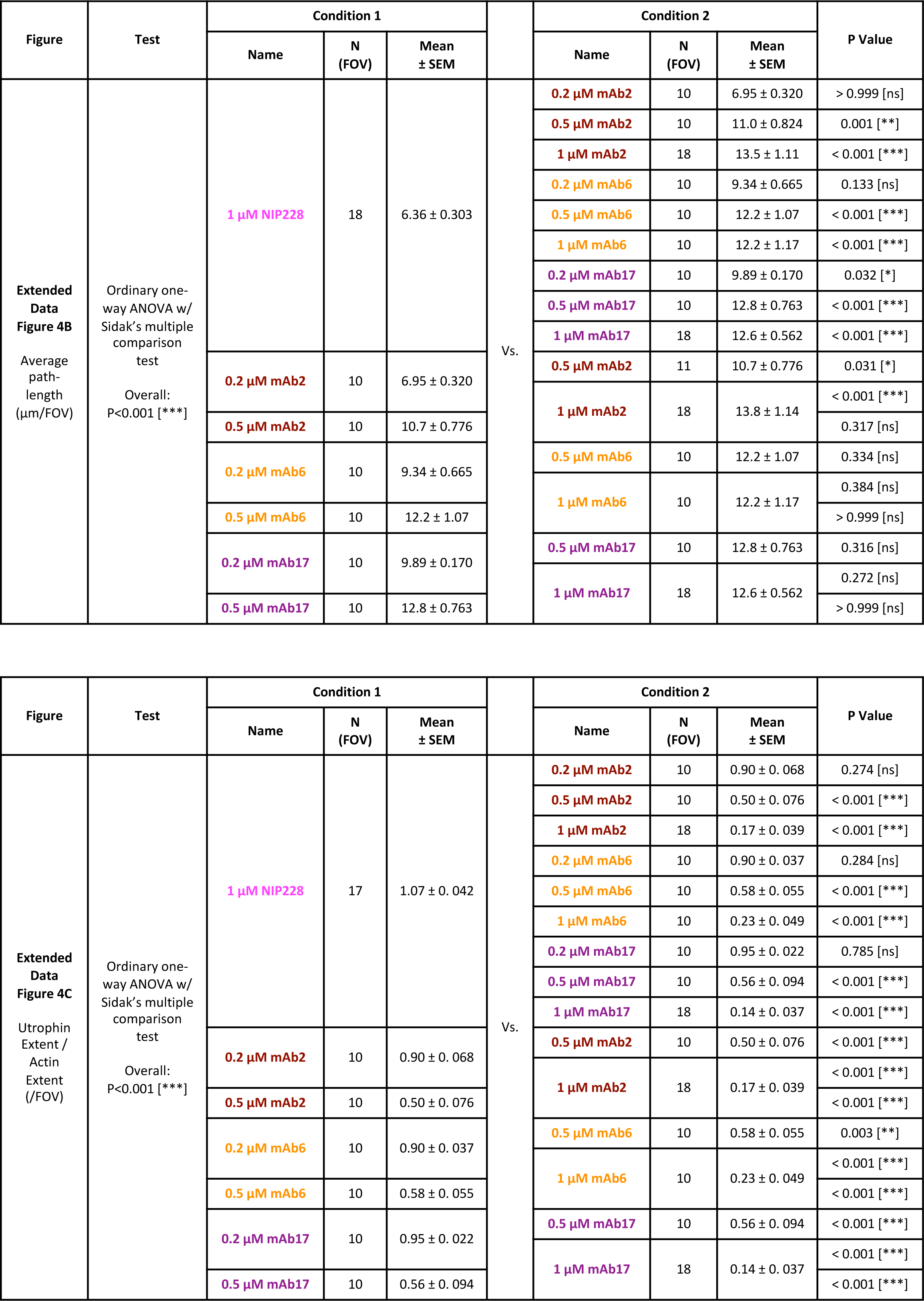
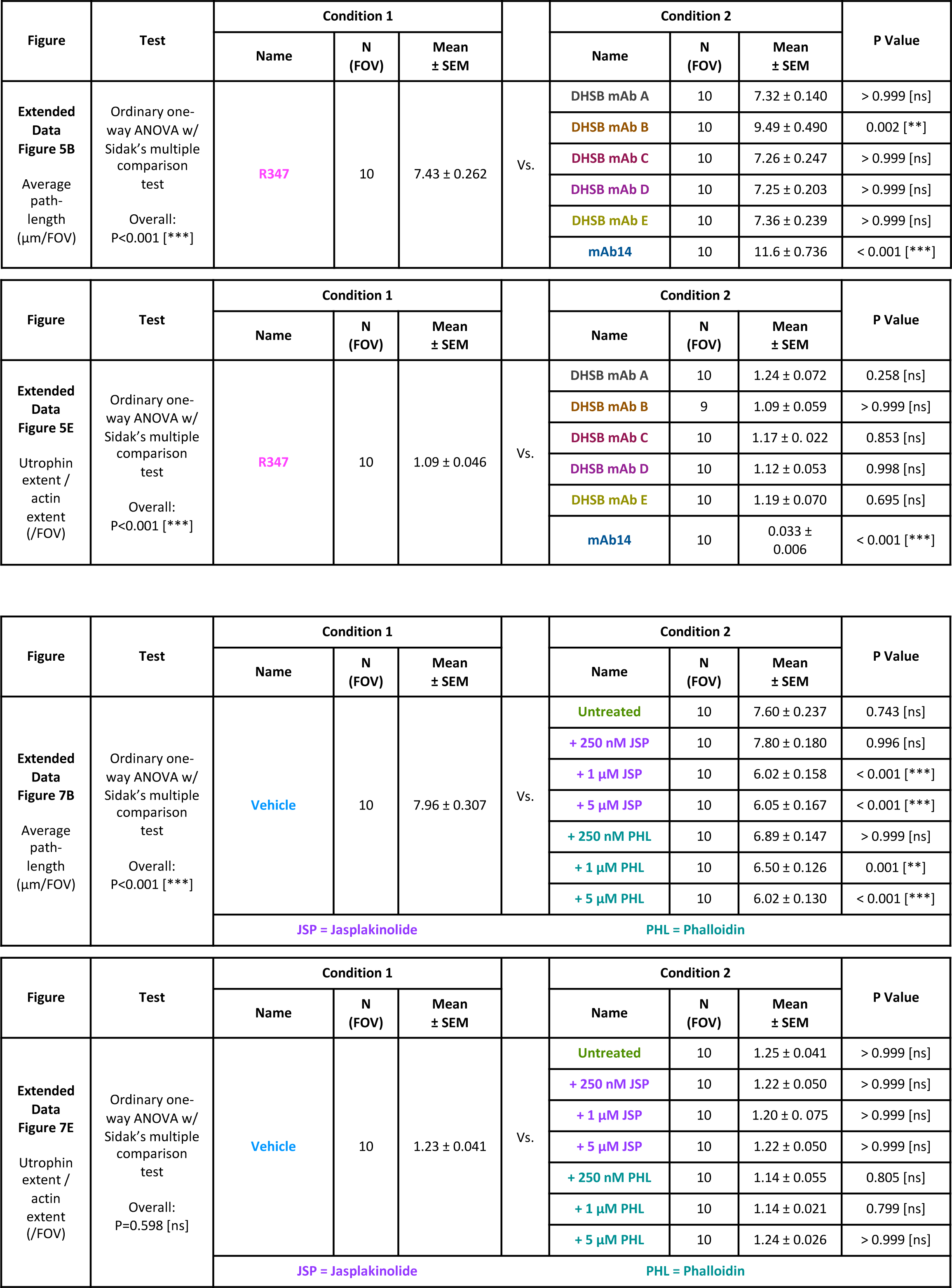
Lists of statistical tests and p values.

## Data and Material Availability

Data is available on request to JLG. The antibodies are available from AstraZeneca with a Material Transfer Agreement.

## Acknowledgments

We thank Thomas Blake, Laura Machesky and Daniel St Johnston for valuable comments on the manuscript. We thank the Gurdon Institute Imaging facility for microscopy support and image analysis. The work was funded by: a Wellcome Trust (www.wellcome.org) Senior Research Fellowship 219482/Z/19/Z to JLG, a MedImmune/AstraZeneca (www.astrazeneca.co.uk) PhD studentship (RG75909/A15360) awarded to CD, JLG and PSI, a BBSRC (www.ukri.org/councils/bbsrc/) Flexible Talent mobility grant RG99217/22799 to JRG. We acknowledge core institute funding by the Wellcome Trust (092096) and Cancer Research UK (www.cancerresearchuk.org/) (C6946/A14492). MedImmune/AstraZeneca scientists were involved in study design, data analysis, decision to publish and preparation of the manuscript and are listed as authors. No other funders played any role in study design, data collection or analysis, decision to publish or preparation of the manuscript.

## Notes

### Competing Interest Statement

The authors have declared no competing interest.

